# Prolonged Lifespan, Ameliorated Cognition, and Improved Host Defense of *Caenorhabditis elegans* by *Lactococcus lactis* subsp. *cremoris*

**DOI:** 10.1101/2021.04.14.439594

**Authors:** Tomomi Komura, Asami Takemoto, Hideki Kosaka, Toshio Suzuki, Yoshikazu Nishikawa

## Abstract

This study evaluated whether the lactic acid bacteria *Lactococcus lactis* subsp. *cremoris* strain FC (FC) could ameliorate host defenses and cognitive ability and extend the lifespan of *Caenorhabditis elegans*, a model of senescence. The lifespan and resistance to physical, chemical, and biological stressors were compared between *C. elegans* fed FC and those fed *Escherichia coli* OP50 (OP), an international standard food for *C. elegans*. Living FC successfully extended the health span, enhanced host defense, and ameliorated the cognitive ability of the nematodes; even the exopolysaccharides (EPSes) of FC could extend the lifespan of *C. elegans*. The chemotaxis index, which was used to evaluate the senescence of sensory neurons, tended to decrease with aging; however, it was more stable in worms fed FC and was significantly higher than that of the control worms at 7 days of age. The worms fed FC were tolerant to *Salmonella enterica* serovar Enteritidis or *Staphylococcus aureus* infection and had better survival than the control worms fed OP. FC showed beneficial effects in *C. elegans daf-16* and *pmk-1* mutants, but not in *skn-1* mutants. Since SKN-1 is the *C. elegans* ortholog of Nrf2, we measured the transcription of heme oxygenase-1 (HO-1), which is regulated by Nrf2, in murine macrophages and found that HO-1 mRNA expression was increased >5 times by inoculation with either FC cells or heat-killed bacteria with EPSes. Thus, both FC and the EPSes can affect longevity *via* the SKN-1/Nrf2 pathway in both nematodes and mammalian cells.

**IMPORTANCE:** Ageing is one of our greatest challenges. The World Health Organization proposed the concept of “Active Ageing” might encourage people to continue to work according to their capacities and preferences as they grow old and would prevent or delay disabilities and chronic diseases that are costly to both individuals and the society, considering that disease prevention is more economical than treatment. Probiotic bacteria such as lactobacilli are living microorganisms that exert beneficial effects on human health when ingested in sufficient amounts and can promote longevity. The significance of this study is that it revealed the anti-senescence and various beneficial effects of the probiotic representative bacterium *Lactococcus lactis* subsp. *cremoris* strain FC and its exopolysaccharides in the nematode *Caenorhabditis elegans*.

## INTRODUCTION

According to the World Health Organization, population ageing is one of humanity’s greatest triumphs, but it can also be our greatest challenge. Population aging has already been recognized in developing countries; by 2025, the number of people aged 60 and over will have increased to approximately 840 million, representing 70% of all older people worldwide. The World Health Organization proposed “Active Ageing: A Policy Framework” that might encourage people to continue to work according to their capacities and preferences as they grow old. Considering that disease prevention is more economical than treatment, this would also prevent or delay disabilities and chronic diseases that are costly to both the individual and the society.

Active aging can be accomplished by consuming healthy foods and nutrients. Based on the longevity of Bulgarians who consumed large quantities of yogurt, Metchnikoff hypothesized that lactobacilli are important for human health and longevity (1). This has accelerated research focusing on healthy bacteria. Probiotic bacteria are defined as living microorganisms that exert beneficial effects on human health when ingested in sufficient amounts (2). The influence of the microbiome on human health is becoming evident, and thus, the dietary inclusion of probiotics, prebiotics, synbiotics, and biogenics is expected to improve health.

*Caenorhabditis elegans* is a small, free-living soil nematode (roundworm) that feeds on bacteria. This nematode has been extensively used as an experimental model for biological studies because of its simplicity, transparency, ease of cultivation, and suitability for genetic analysis (3). Furthermore, due to its short and reproducible life span, *C. elegans* is particularly suitable for aging studies (4). One century after Metchnikoff’s study, we succeeded in showing that lactic acid bacteria (LAB) exert longevity effects in *C. elegans* (5). At present, the *C. elegans* model is being used worldwide to evaluate probiotic strains (6, 7), and the strains selected using this model have shown beneficial effects even in swine (8).

LAB have been used in various fermented foods since antiquity and are the most used probiotic microorganisms. The LAB sub-specie, *Lactococcus lactis* subsp. *cremoris*, is expected to have various physiological effects on humans consuming probiotic dairy foods. We have reported various beneficial effects of *L. lactis* subsp. *cremoris* strain FC (FC) on colitis (9), immunomodulation (10), biopreservation, antimicrobial activities (11), dermatitis (12), and healthy defecation (13). Furthermore, other groups reported the benefits of *L. lactis* for folate supplementation (14) and its inhibitory effects on food allergies (15), pollinosis (16), tumors (17), serum cholesterol (18), and hypertension (19). In addition, strain H61 was shown to suppress aging-associated symptoms such as osteoporosis (20) and hearing loss (21) in senescence-accelerated mice.

In this study, we evaluated whether strain FC could ameliorate host defenses, retard the accumulation of advanced glycation end-products (AGEs) and deterioration of cognitive ability, and prolong the lifespan of *C. elegans*. The life span and resistance to physical, chemical, and biological stressors were compared between *C. elegans* fed FC and those fed *Escherichia coli* OP50 (OP), an international standard food for *C. elegans.* In addition, the mechanism of the effects of FC feeding were determined using *C. elegans* mutants, in which defense-related genes were unfunctional. Human macrophages were also inoculated with FC to investigate whether the bacteria could influence human innate immunity.

## RESULTS

### Amelioration of senescence

The average life span of the nematodes fed FC was greater than that of nematodes fed OP, irrespective of whether the FC organisms were cultured aerobically or anaerobically (Fig. 1A). Both living and heat-killed FC prolonged the lifespan of *C. elegans*, compared to the nematodes fed heat-killed OP. In contrast, the life span of *C. elegans* fed heat-killed OP was longer than that of *C. elegans* fed living OP because of the loss of virulence (Fig. 1B). The longevity afforded by FC was not attributed to its non-pathogenic properties alone.

**Fig. 1A.**
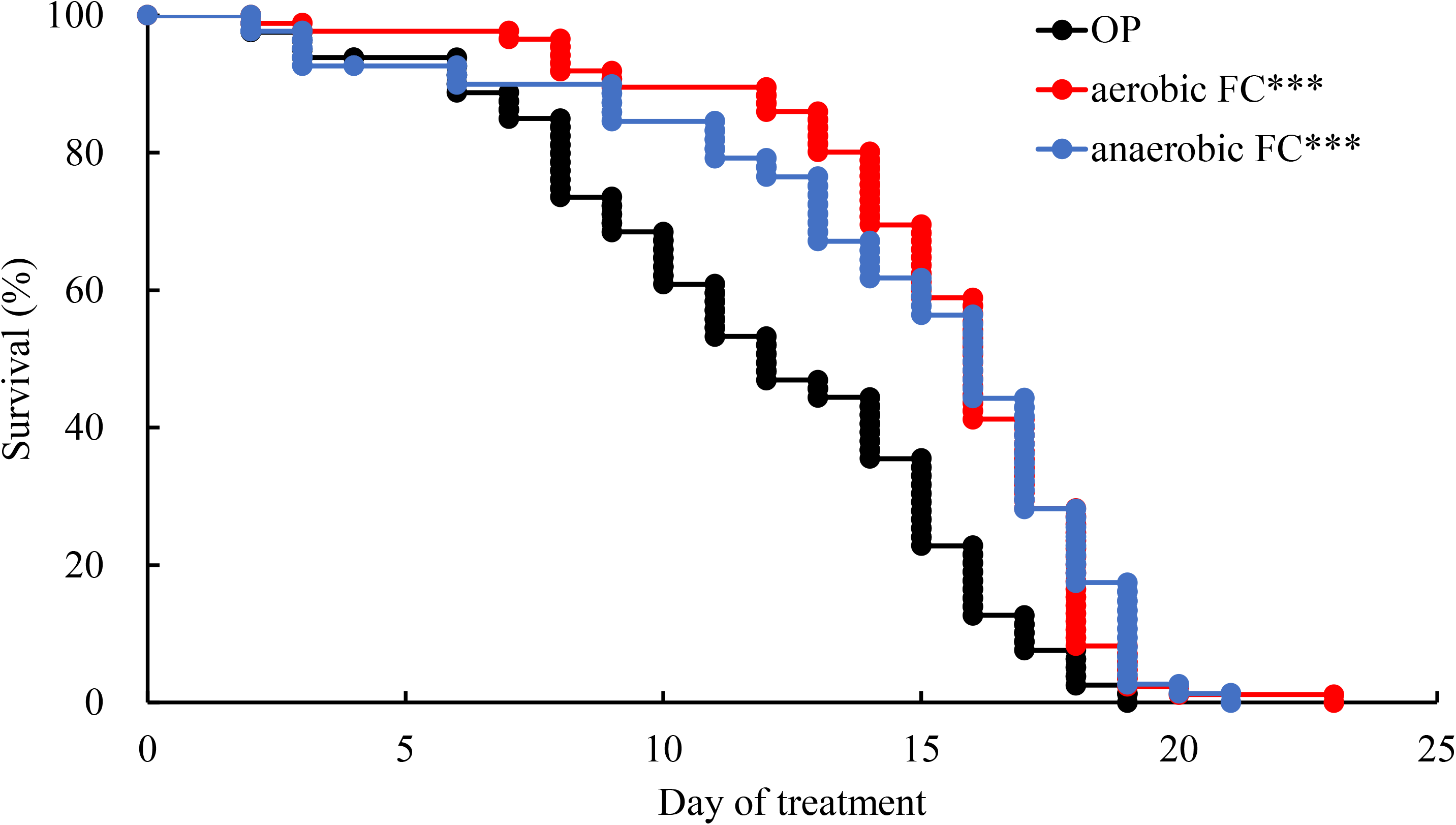
Anti-senescence effects of *L. lactis* subsp. *cremoris* FC (FC) on *C. elegans*. Survival curves of *C. elegans* fed FC cultured under aerobic or anaerobic conditions compared with the lifespan of control worms fed only a diet of *E. coli* strain OP50 (OP). Each plate contained 10 mg (wet weight) of bacteria. Worms were 3 days old on day 0. The lifespan of the worms fed FC (both aerobic and anaerobic conditions) were longer than that of worms fed OP. Each assay was repeated five times by using aerobically cultured organisms and once with anaerobically cultured organisms. Table 2 summarizes the data obtained from all experiments, and the curves were drawn based on representative experiments. Asterisks indicate significant differences (*** *p* < 0.001) from control worms fed OP.

**Fig. 1B.**
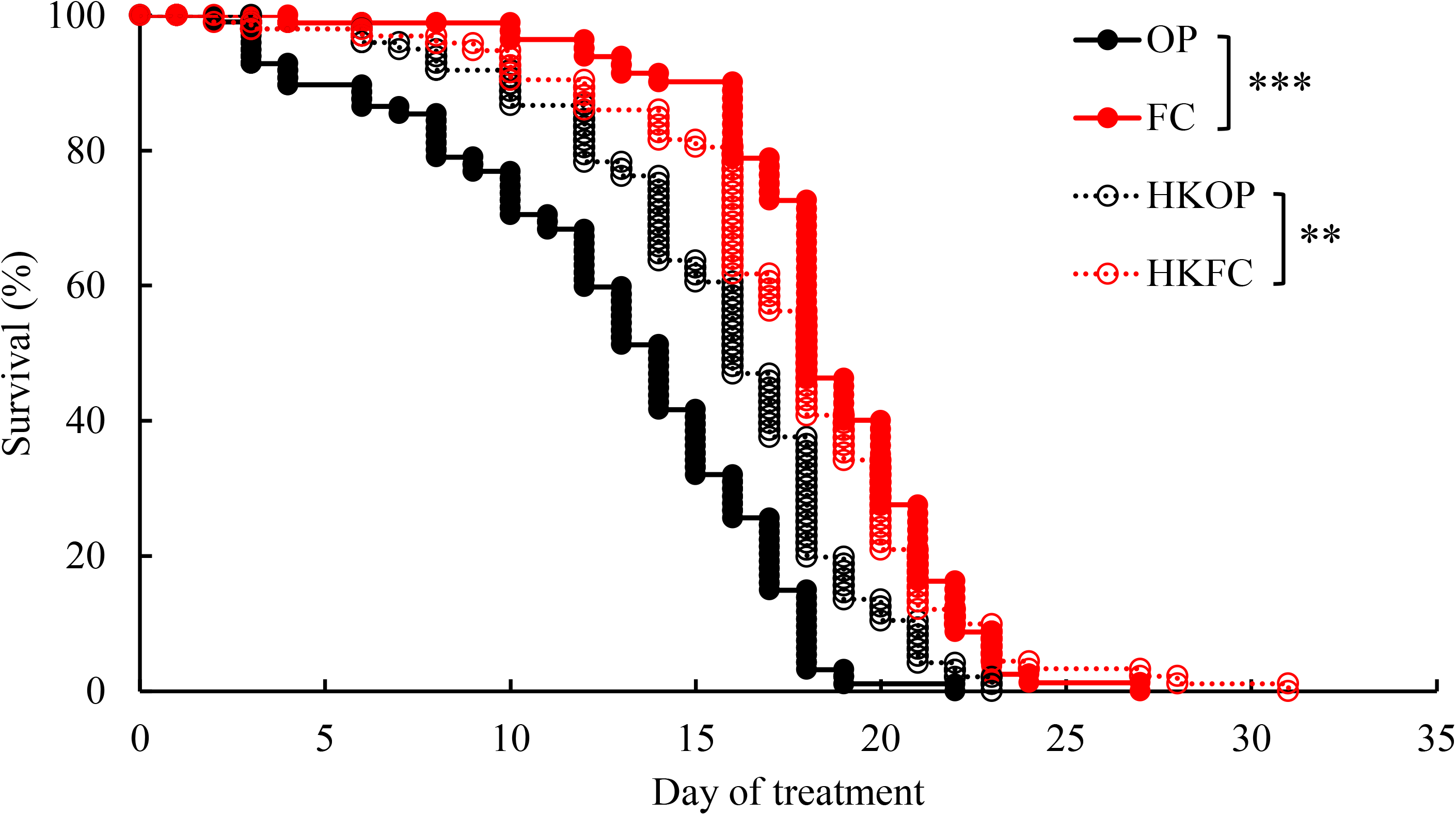
Anti-senescence effects of FC on *C. elegans*. Survival curves of *C. elegans* fed heat-killed bacteria. Adult worms fed OP for 3 days after hatching were placed onto plates containing heat-killed (HKOP, HKFC) or living bacteria (OP, FC). Worms were 3 days old at day 0. The lifespan of worms fed heat-killed OP was shorter than that of the worms fed living or heat-killed FC. Each assay was repeated three times. Table 2 summarizes the data obtained from all experiments, and the curves were drawn based on representative experiments. Asterisks indicate significant differences (** *p* < 0.01, *** *p* < 0.001) from control worms fed living OP or heat-killed OP.

**Fig. 1C.**
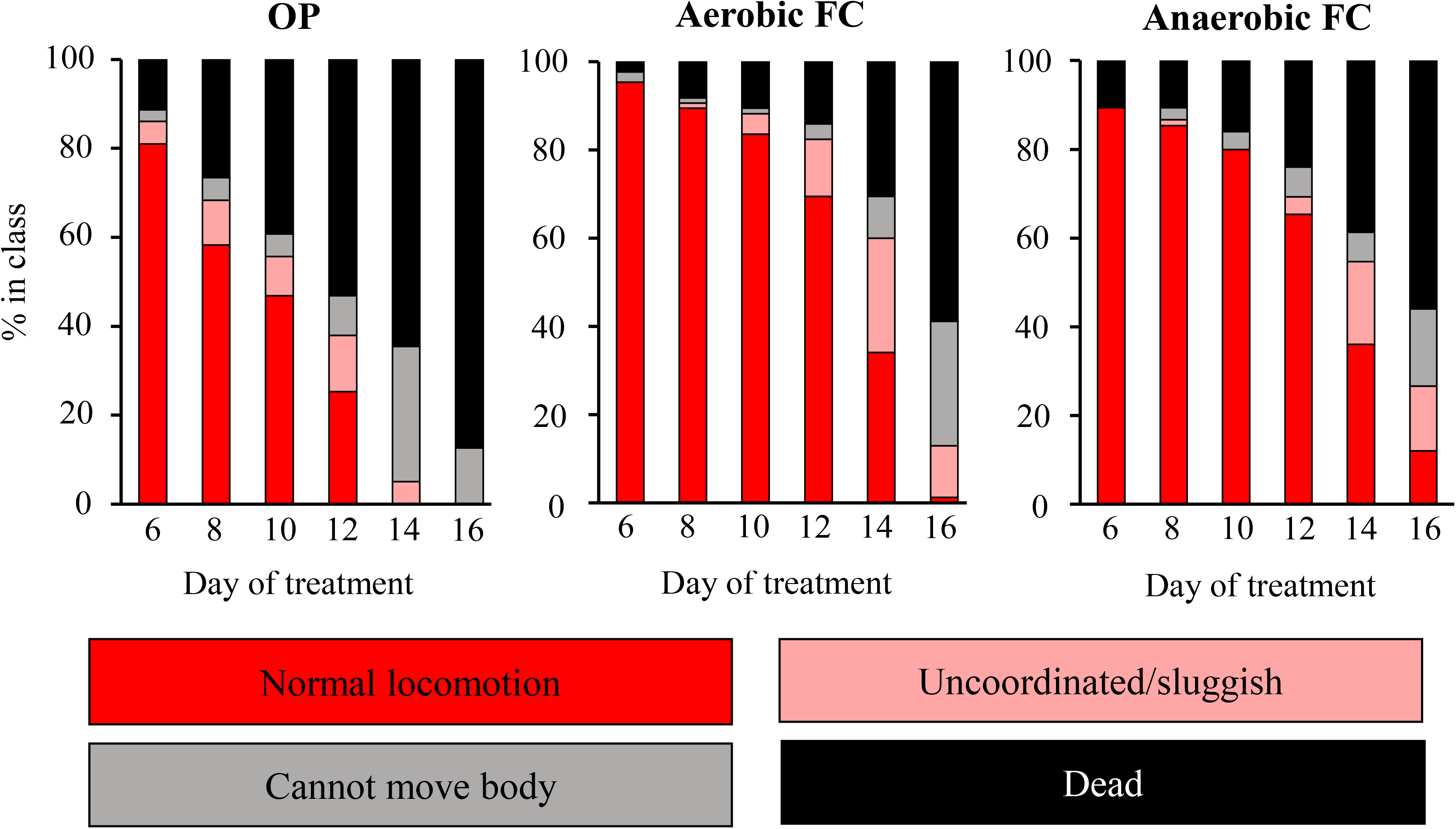
Anti-senescence effects of FC on *C. elegans*. Locomotory activity of *C. elegans* fed FC. Young adult worms fed OP for 2 days after hatching were transferred to plates containing 10 mg of either OP or FC cultured aerobically or anaerobically on the surface. Worms were 3 days old on day 0. Animals were classified into four classes based on their locomotion: class A, robust, coordinated sinusoidal locomotion (red bars); class B, uncoordinated and/or sluggish movement (pink bars); class C, no forward or backward movement, but head movements or shuddering in response to prodding (gray bars); and class D, dead animals (black bars). The proportion of each class at the indicated time point are indicated. The proportion of class A was greater in both the FC groups than in the control group fed OP. Each assay was repeated twice by using aerobically cultured organisms and once with anaerobically cultured organisms.

**Fig. 1D.**
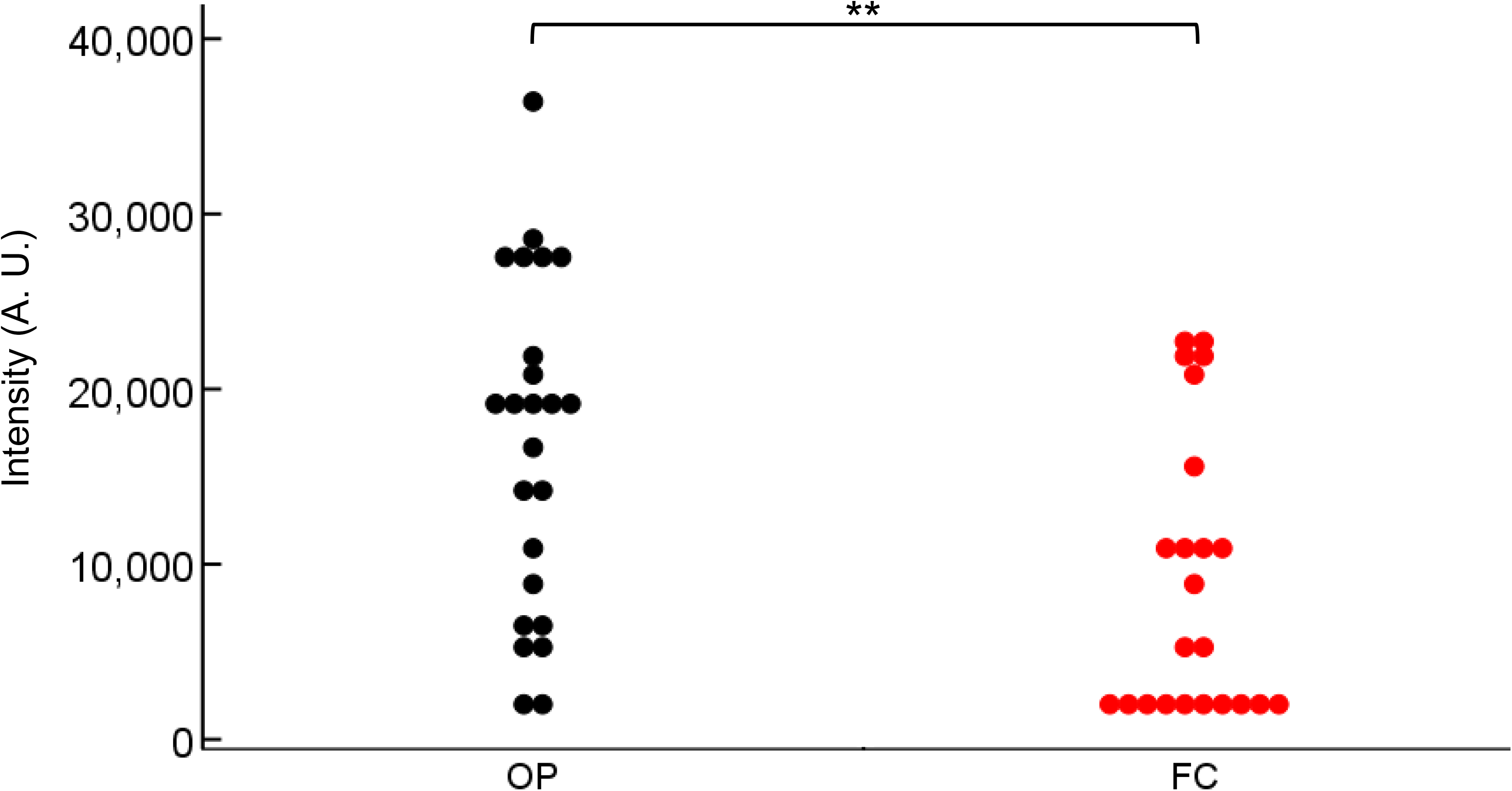
Anti-senescence effects of FC on *C. elegans*. Autofluorescence (ex. 340 nm/430 nm) in 9-day-old worms grown with OP or aerobic FC was measured using the non-invasive wrap-drop method. Randomly selected worms were washed with M9 buffer and then placed in 1.0 µL M9 buffer on a cling film stretched over a 384-well black plate. The autofluorescence in the body of each worm was captured using the multimode grating microplate reader. Intensity of blue autofluorescence in the FC-fed worms was weaker than that in the control worms fed OP. Each assay was performed with a minimum of 20 worms. Asterisks indicate statistically significant difference (** *p* < 0.01) from control worms fed OP.

**Fig. 1E.**
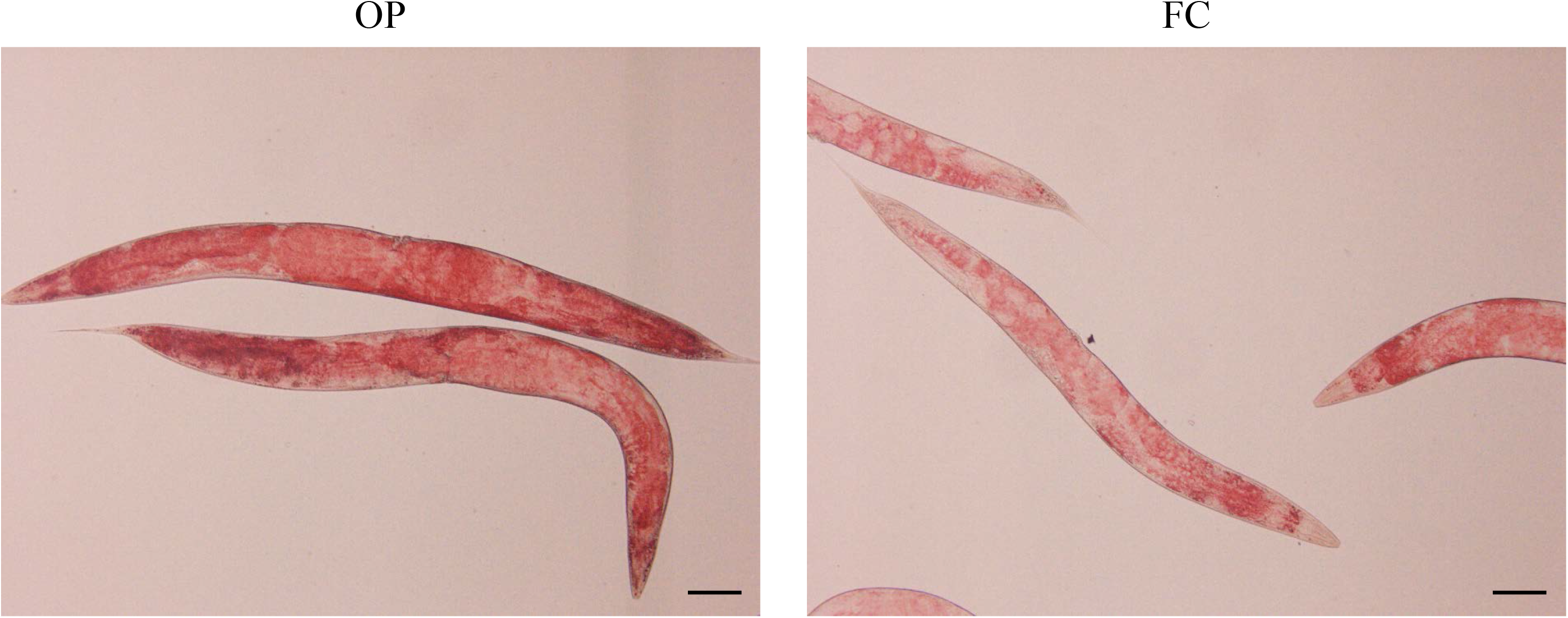
Anti-senescence effects of *L. lactis* subsp. *cremoris* FC on *C. elegans*. Lipid accumulation in nematodes. Oil red O staining in age-synchronized worms fed OP was apparent in 7-day-old worms but weaker in worms fed aerobic FC. The assay was repeated twice. Scale bar indicates 100 μm.

**Fig. 1F.**
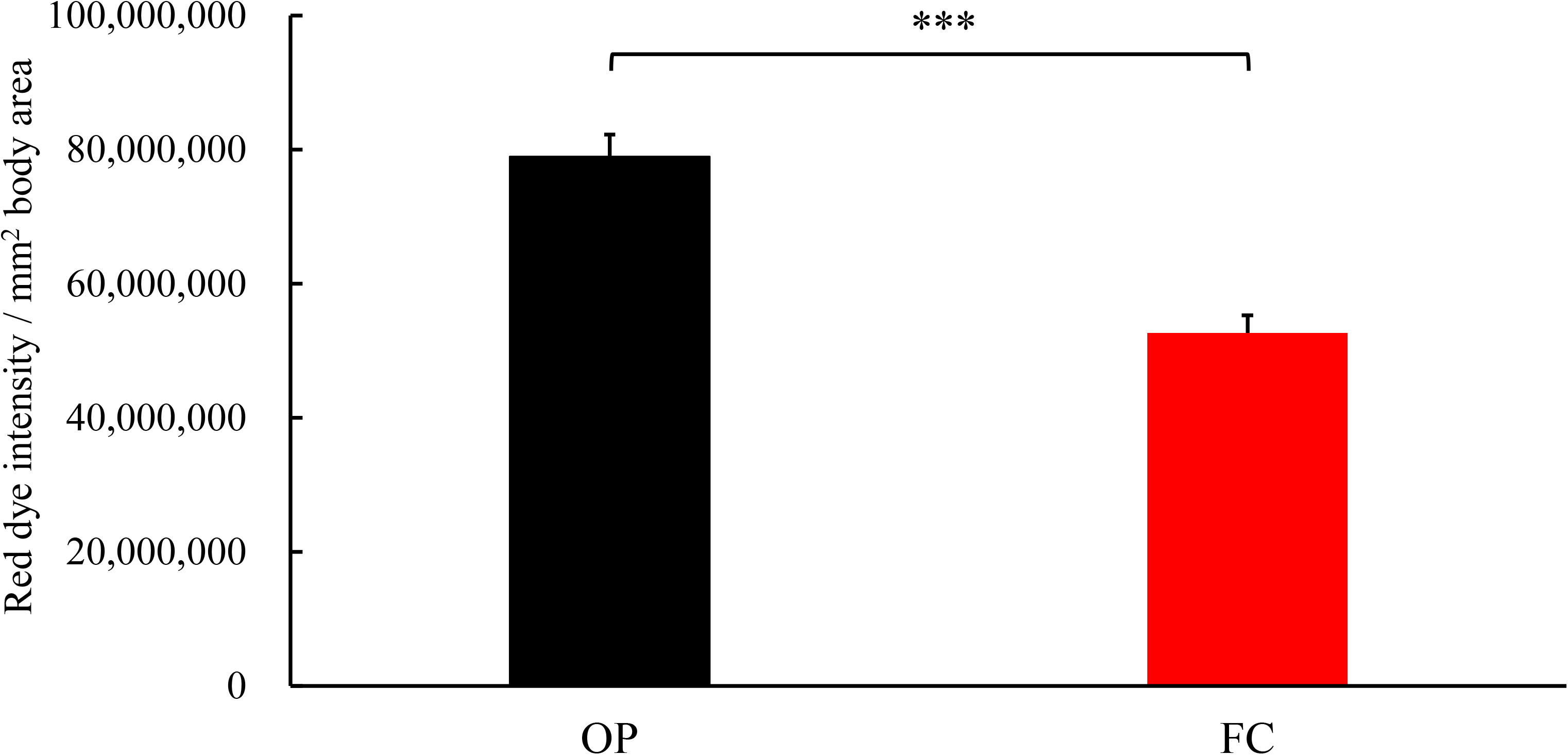
Anti-senescence effects of FC on *C. elegans.* Red dye on 7-day-old worms was quantified based on the projection area of a worm’s body. Each bar represents the average values for ten worms. Error bars represent the standard error. Asterisks indicate statistically significant difference (*** *p* < 0.001) from control worms fed OP.

**Fig. 1G.**
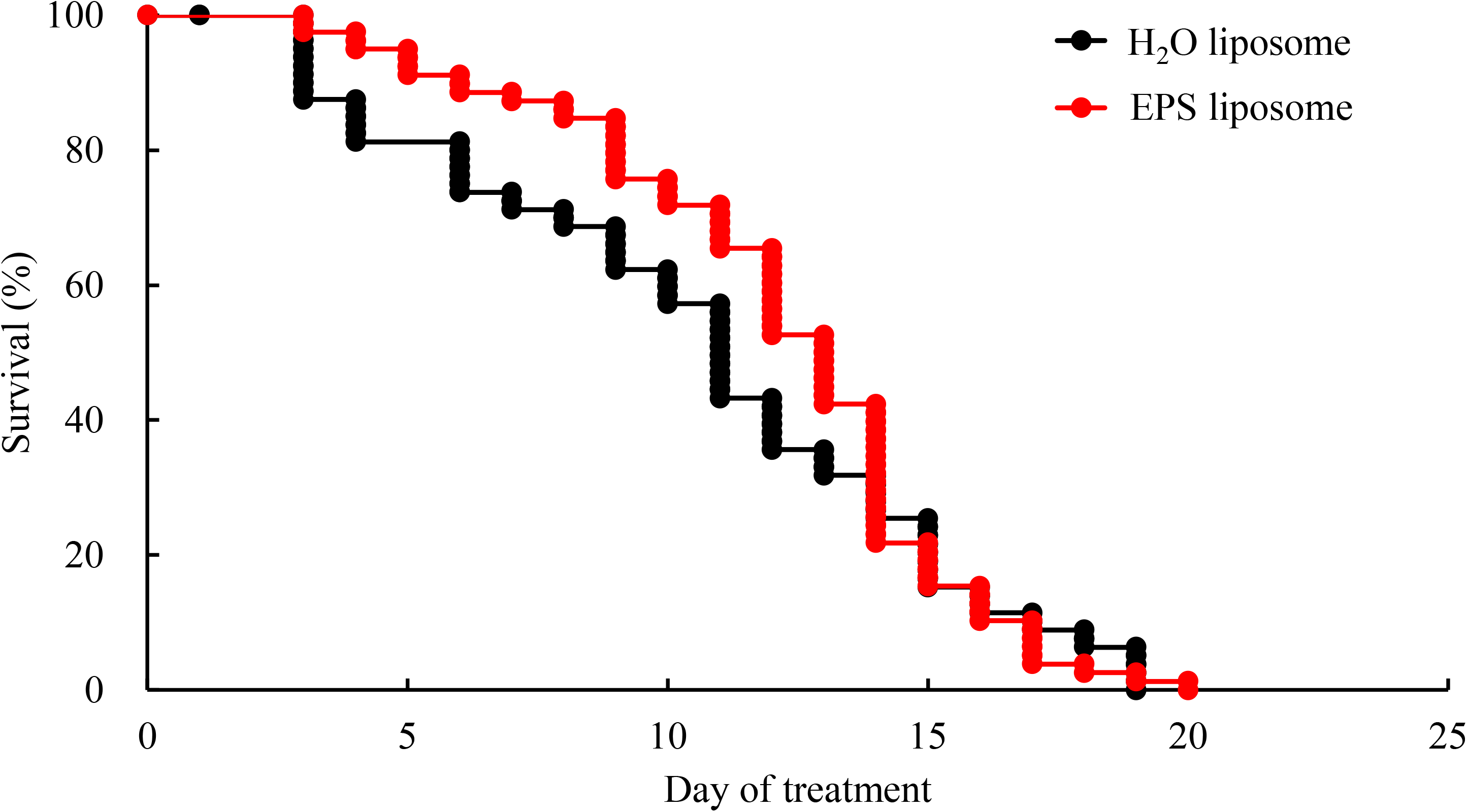
Anti-senescence effects of FC on *C. elegans.* Survival curves of *C. elegans* fed the exopolysaccharide (EPS) extracted from FC. Adult worms fed OP for 3 days after hatching were placed onto plates containing OP (10 mg) mixed with the EPS liposome or H_2_O liposome as control. Worms were 3 days old at day 0. The lifespan of worms fed EPS liposome was slightly prolonged compared to those fed H_2_O liposomes. Each assay was repeated three times. Table 2 summarizes the data obtained from all experiments, and the curves was drawn based on representative experiments.

**Table 1.**
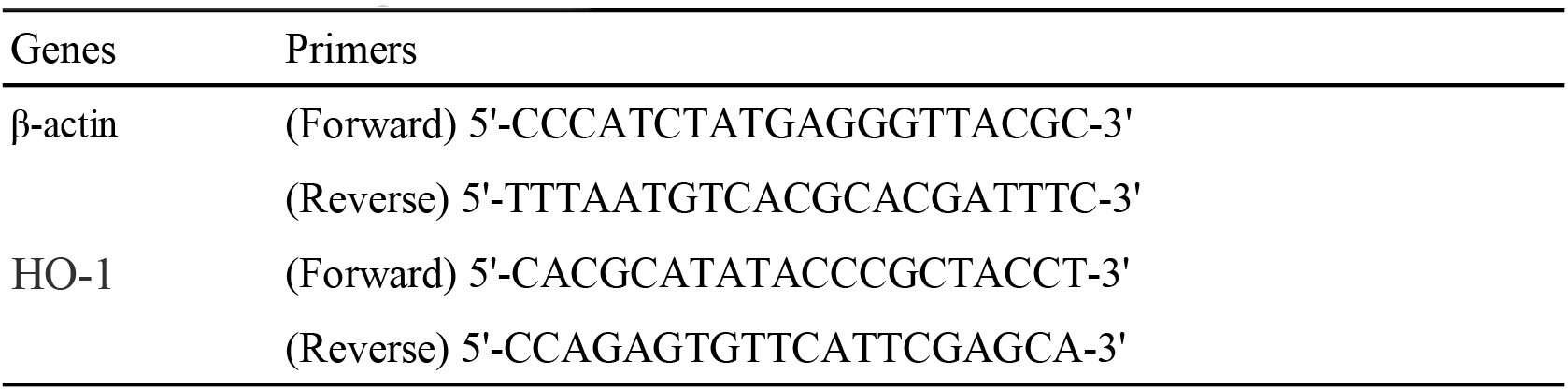
Primers for qPCR

**Table 2.**
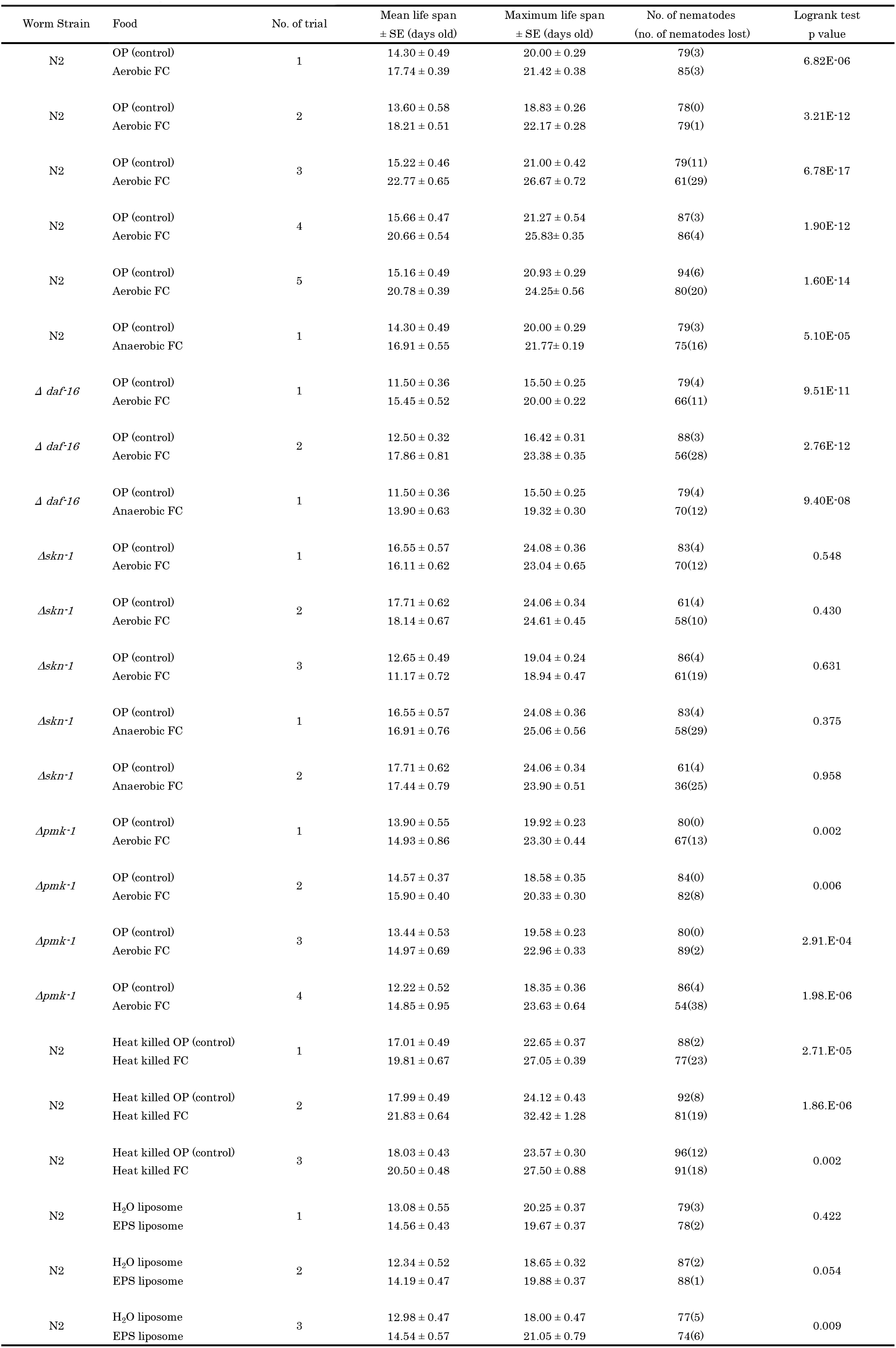
Mean and maximum life span of worms fed FC or EPS

Locomotor ability was measured as an indicator of health span. The ratio of nematodes showing coordinated sinusoidal locomotion (class A) was higher in the FC-fed group than in the control nematodes fed OP (Fig. 1C).

Recently, we showed that autofluorescence was greater in older *C. elegans* than in young ones, and this was possibly attributed to the AGEs of vitellogenins (22). Since FC prolonged the lifespan of nematodes, FC-fed nematodes were assumed to be less autofluorescent. Indeed, the intensity of blue autofluorescence in FC-fed *C. elegans* was weaker than that in the control *C. elegans* fed OP (Fig. 1D). The effects of FC on the metabolism of *C. elegans* were also determined to assess how FC prolonged the life span of the nematodes. Oil Red O staining showed that the nematodes fed FC stored less lipids (Fig. 1E and F).

The production of exopolysaccharide (EPS) is a well-known characteristic of *L. lactis* subsp. *cremoris* (23). As the longevity effect of heat-killed FC was somewhat lower than that of living FC, we attempted to determine whether the longevity effect was attributed to EPSes purified from FC. By using our previously reported liposome method (24), liposomes, including acellular EPSes, were administered to the nematodes with OP. Apart from FC cells, EPSes independently tended to prolong the lifespan of the nematodes and possibly showed anti-senescence effects (Fig. 1G).

As FC extended the health span of *C. elegans*, we investigated whether the bacteria could ameliorate the senescence of the cognitive ability of *C. elegans* (Fig. 2A). The chemotaxis index, which tends to decrease with aging, was more stable and significantly higher in the nematodes fed FC than in the control nematodes at 7 days of age (Fig. 2B).

**Fig. 2A.**
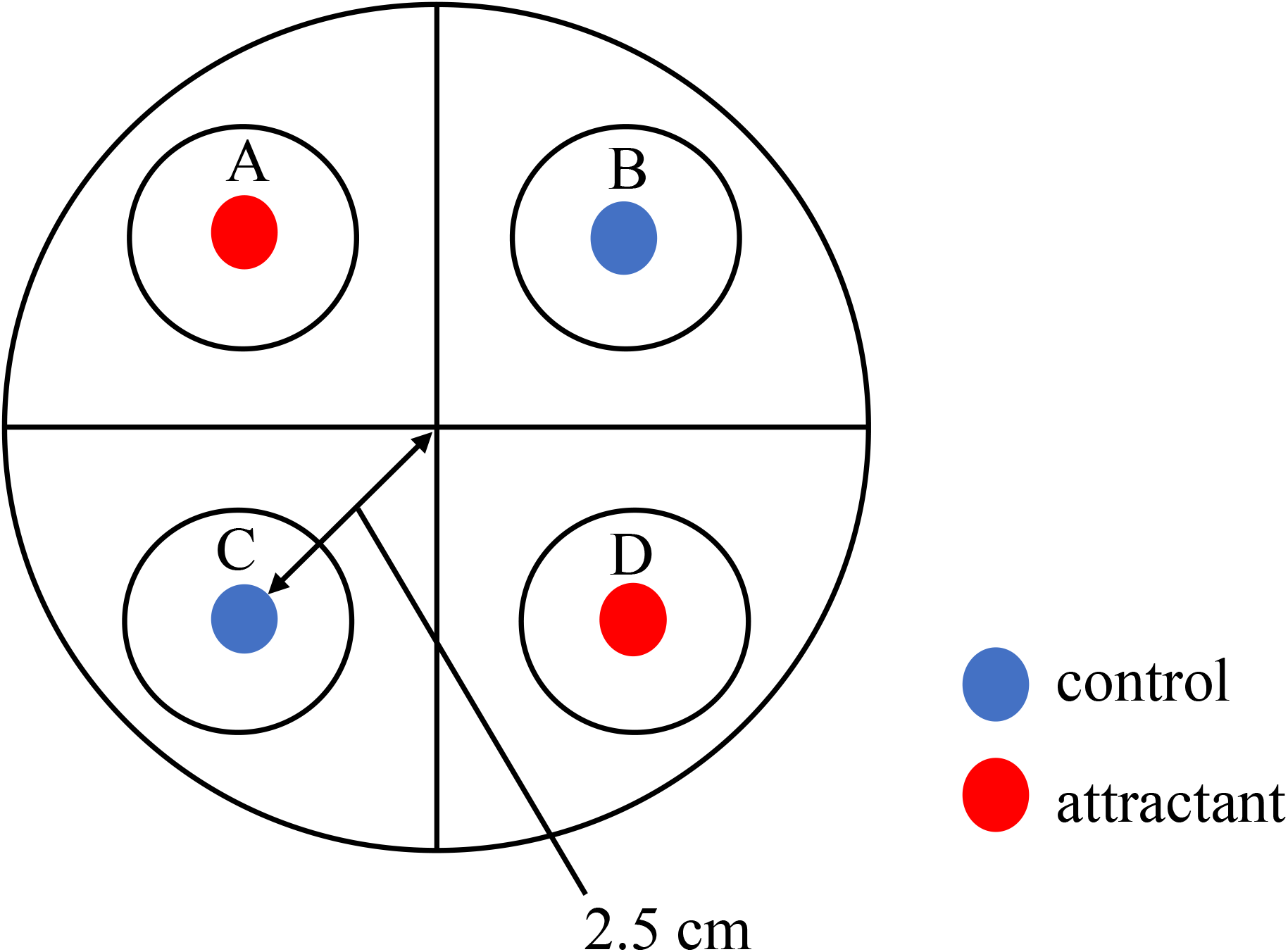
Effects of FC on the cognitive ability of *C. elegans*. Schematic view of the chemotaxis assay. The wild-type N2 strain was fed with OP or aerobic FC from 3 days of age. Assay plates were divided into quadrants of two attractants (A & D) and two controls (B & C). Sodium azide was also included with the attractant and control to paralyze the worms. The worms were transferred to the center of the assay plate and incubated at 25°C for 2 h. Next, the number of worms in each quadrant was scored. The chemotactic index (CI) was calculated using the formula: CI = (number of worms in both attractant quadrants A and D – number of worms in both control quadrants B and C)/total number of worms on the assay plate.

**Fig. 2B.**
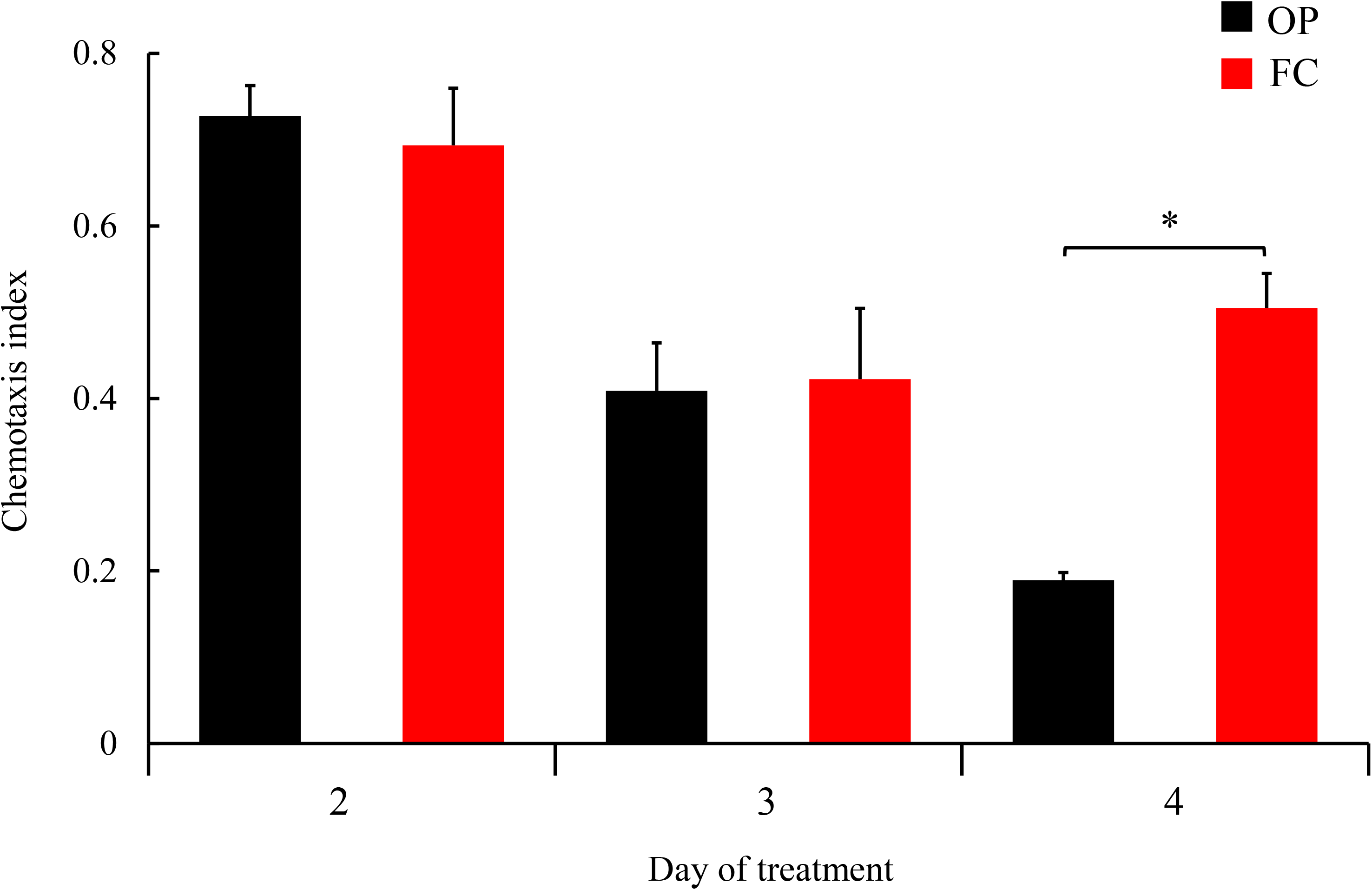
Effects of FC on the cognitive ability of *C. elegans*. The CI of wild-type N2 strain fed OP or aerobic FC from 3 days of age. The CI of worms fed FC was significantly higher than that of control worms at 7 days of age. Each assay was performed in triplicate and repeated three times. Each bar represents the average values of a minimum of 200 worms. Error bars represent the standard error. Asterisks indicate significant differences at * *p* < 0.05.

We have shown that lactobacilli and bifidobacteria could increase the tolerance of the nematodes to *Salmonella* infection (5). In this study, we investigated whether FC could enhance the host defense of *C. elegans* to *Salmonella enterica* serovar Enteritidis and *Staphylococcus aureus* infection. FC-fed nematodes had better survival than the OP-fed ones after the infection with pathogens (Fig. 3A); this effect was unlikely to be attributed to the increased resistance, as no significant difference in the number of pathogens recovered from both FC- and OP-fed nematodes was noted (Fig. 3B). Zhao et al. (25) reported that pathogens that are virulent to nematodes can invade the oropharyngeal tissues. Indeed, the oropharyngeal epithelia of the nematodes were leaky after infection with *S.* Enteritidis or *S*. *aureus* (Fig. 3C–E); however, the barrier function of the epithelia in FC-fed nematodes made them resistant to pathogen destruction.

**Fig. 3A.**
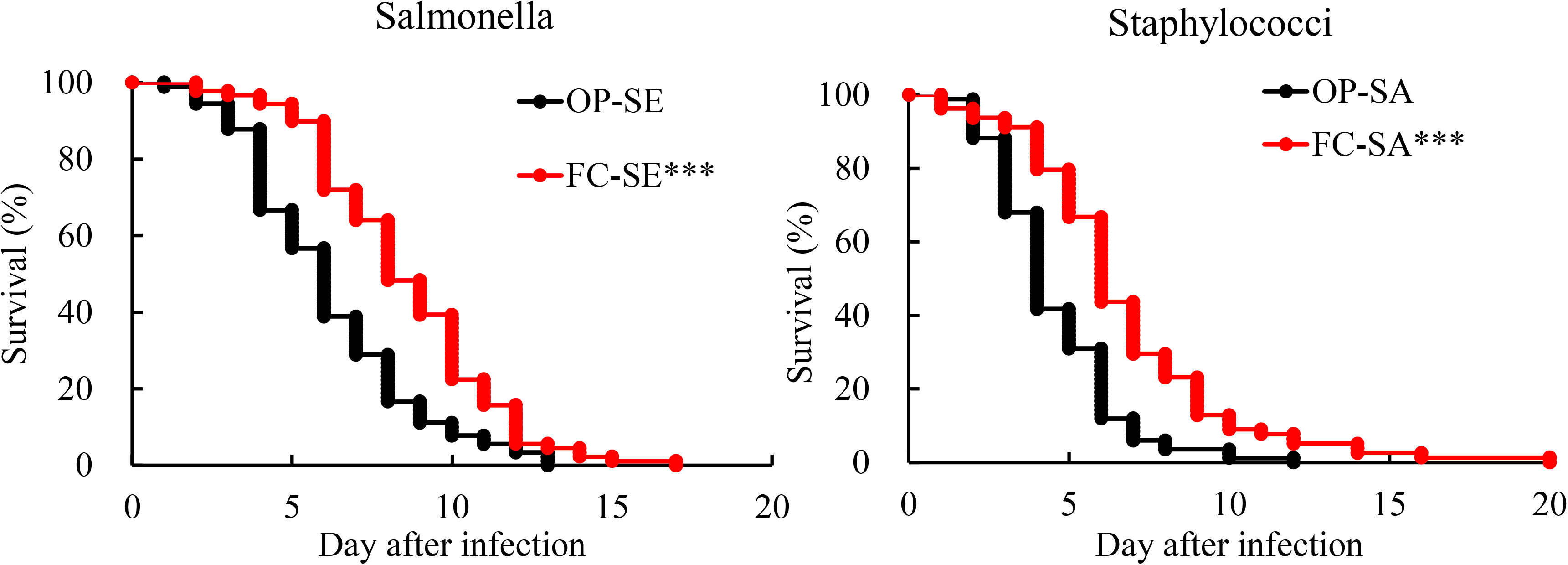
Effects of FC on host defense of *C. elegans*. Effects of FC on the tolerance of the worms to *Salmonella enterica* serovar Enteritidis (SE) or *Staphylococcus aureus* (SA) infection. Adult worms fed OP for 3 days after hatching were transferred to a diet of aerobically grown FC or OP as a control. After 4 days, the nematodes were transferred to SE or SA plates, and survival curves were drawn. Nematodes fed FC were significantly more resistant than controls to the pathogens. Asterisks indicate significant differences (*** *p* < 0.001) from control worms fed OP. Table 3 summarizes the data obtained from all experiments, and the curves were drawn based on representative experiments. Each assay was repeated three times.

**Fig. 3B.**
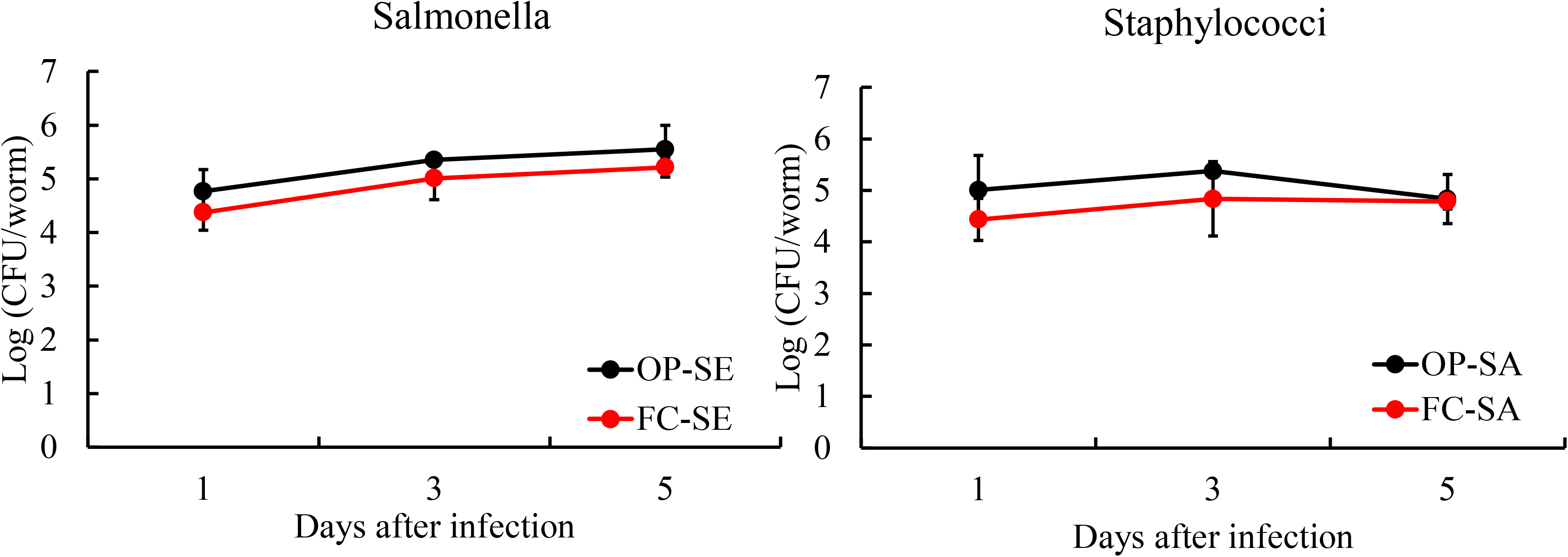
Effects of FC on host defense of *C. elegans*. The colony forming units (CFUs) of SE or SA cells recovered from infected nematodes were obtained by plating whole lysates of 8-, 10-, and 12-day-old worms (1, 3, and 5 days after the start of pathogen ingestion, respectively). The numbers of each pathogen were similar in both groups fed either FC or OP. Each assay was performed on 5 worms and examined once with SE and twice with SA.

**Fig. 3C.**
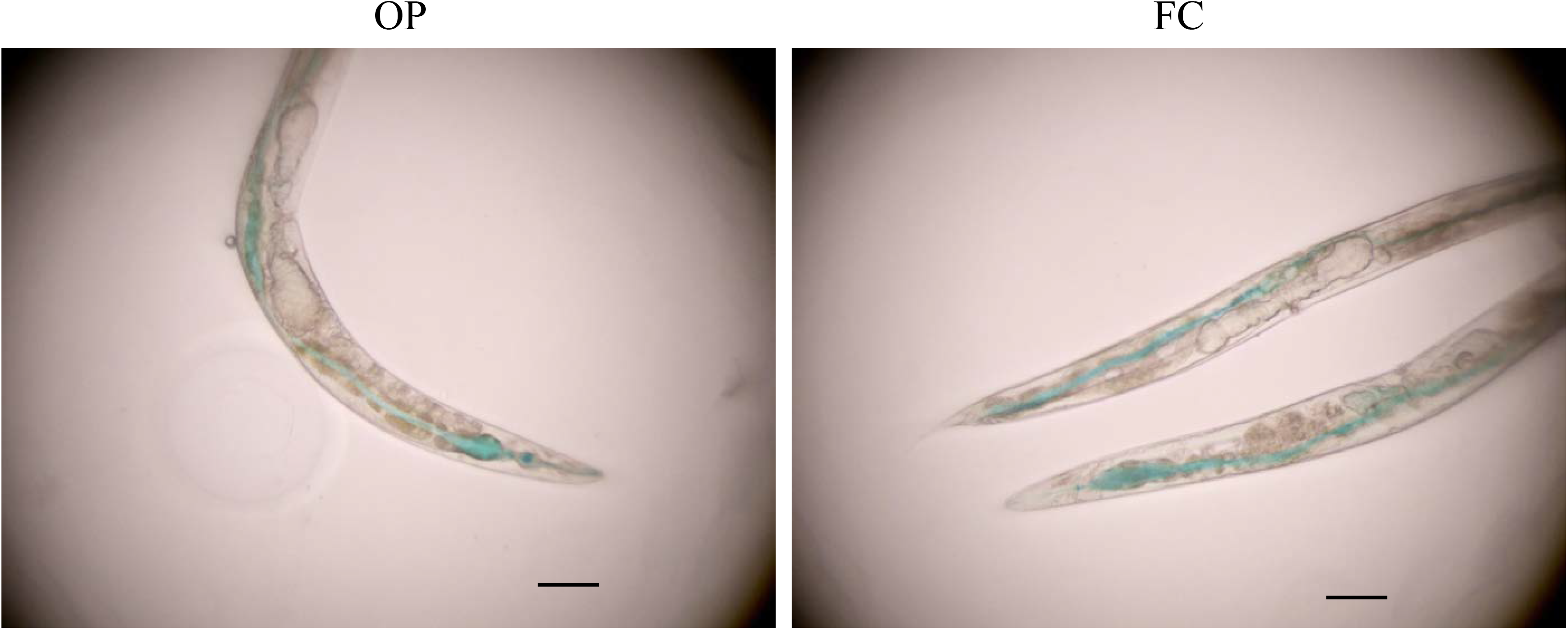
Effects of FC on host defense of *C. elegans*. The intestinal barrier function by using the Smurf assay. Adult worms fed OP for 3 days after hatching were transferred to the diet of aerobically grown FC or OP as a control, and then soaked in blue dye for 3 h at 12 days of age. The scale bar indicates 100 μm.

**Fig. 3D.**
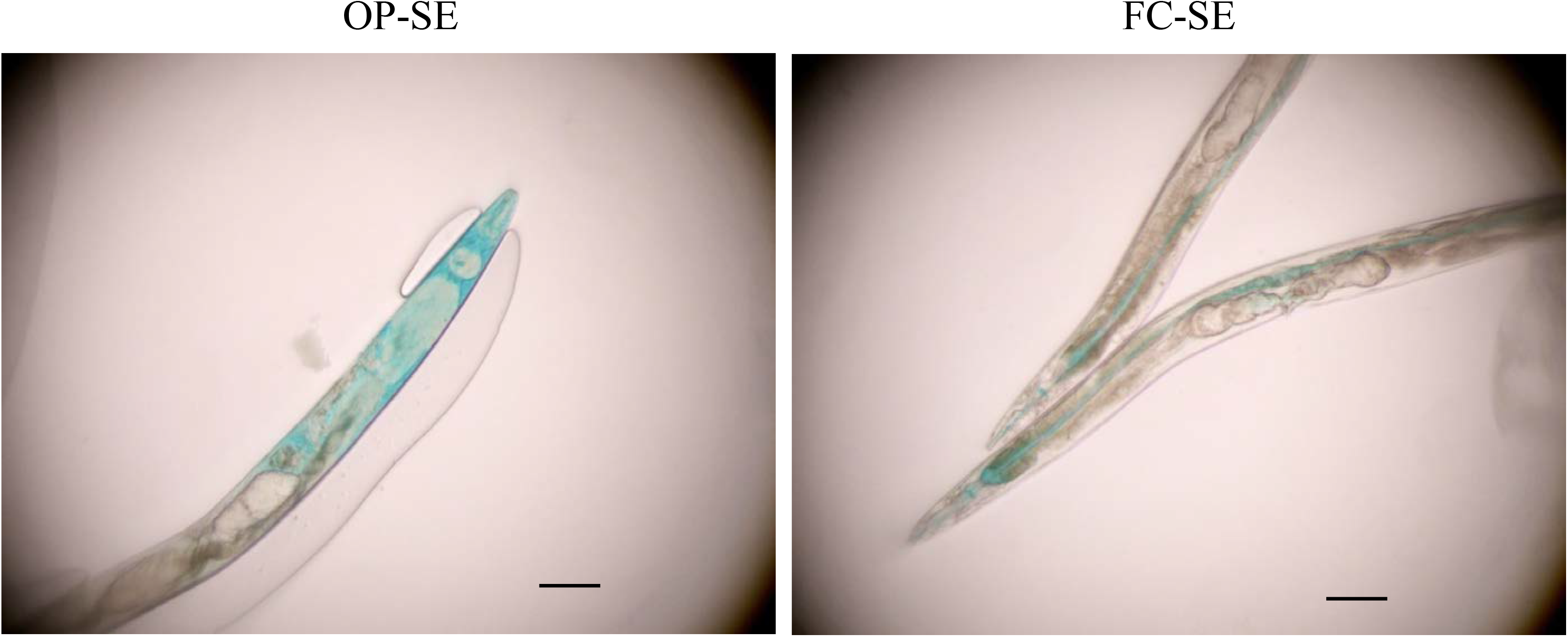
Effects of FC on host defense of *C. elegans*. The intestinal barrier function by using the Smurf assay. After feeding on FC or OP for 4 days, the 7-day-old nematodes were infected with SE, and then stained with blue dye at 12 days of age. In contrast to the differential interference contrast (DIC) images of worms fed OP or aerobically grown FC from 3 days to 12 days without infection, worms infected with pathogens showed spreading of the dye into their body cavity (OP-SE). However, FC-fed worms counteracted the dye spreading to the body cavity, which supports its protective function in the worm gut (FC-SE). The assay was performed twice. Each scale bar indicates 100 μm.

**Fig. 3E.**
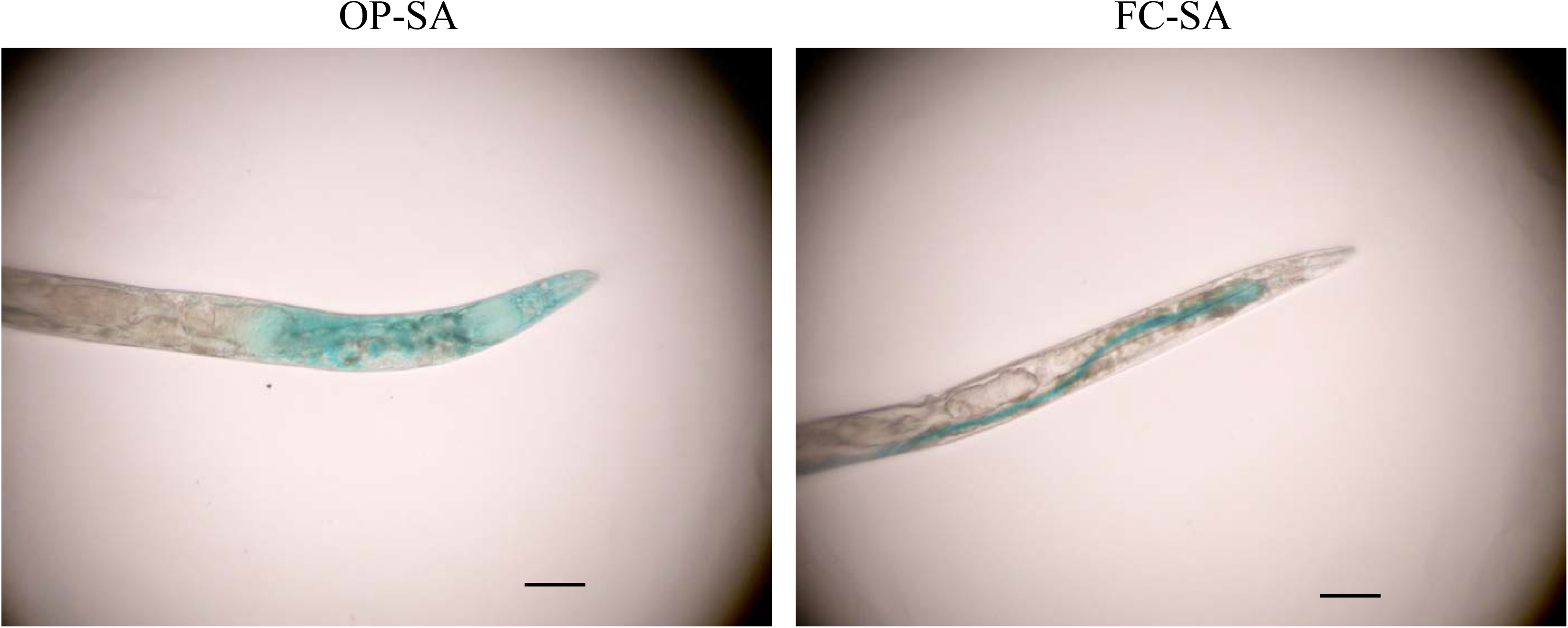
Effects of FC on host defense of *C. elegans*. The intestinal barrier function by using the Smurf assay. After feeding on FC or OP for 4 days, the 7-day-old nematodes were infected with SA, and then stained with blue dye at 12 days of age. In contrast to the DIC images of worms fed OP or aerobically grown FC from 3 days to 12 days without infection, worms infected with pathogens showed spreading of the dye into their body cavity (OP-SA). However, FC-fed worms counteracted the dye spreading to the body cavity, which supports its protective function in the worm gut (FC-SA). The assay was performed twice. Each scale bar indicates 100 μm.

**Fig. 3F.**
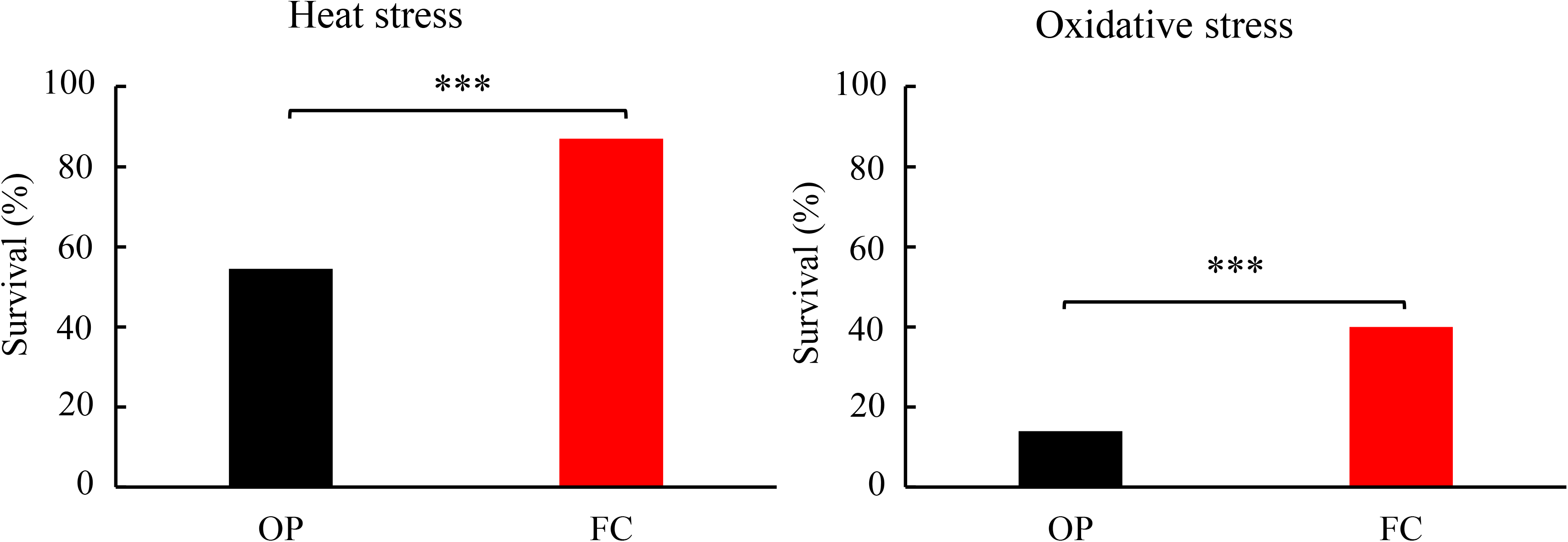
Effects of FC on host defense of *C. elegans*. Influence of FC feeding on the susceptibility of worms to heat and oxidative stress caused by juglone. Worms were fed aerobically grown FC for 4 days beginning from 3 days of age and then either incubated at 35°C for 7 h or in 250 µM juglone for 2 h. The assay was performed with 100 worms and repeated thrice. Asterisks indicate statistically significant differences (*** *p* < 0.001) compared to the control worms exposed to either heat or juglone, respectively.

**Table 3.**
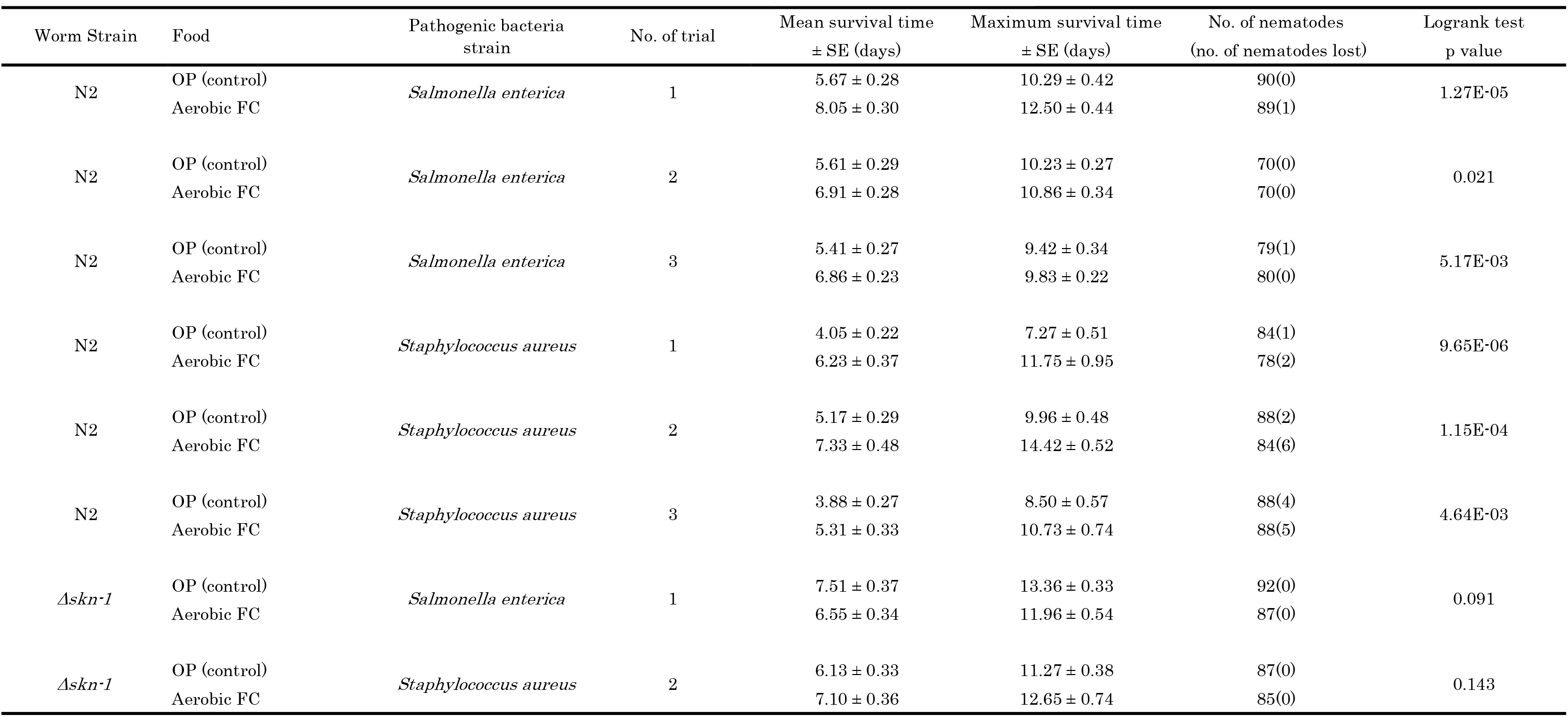
Mean and maximum survival days of worms infected with *Salmonella enterica* serovar Enteritidis or *Staphylococcus aureus*

FC-fed *C. elegans* were resilient against not only senescence and infectious diseases but also physical or chemical stresses. The survival of FC-fed nematodes was better than that of OP-fed ones after exposure to either heat at 37°C or juglone (Fig. 3F); however, FC-fed nematodes could not live significantly longer than the OP-fed ones after exposure to cupric chloride or paraquat (data not shown).

### Mechanisms of anti-senescence by FC

Caloric restriction (dietary restriction in the case of *C. elegans*) is generally considered to extend the lifespan of various animals. The body size of the nematodes fed FC was slightly smaller than that of OP-fed nematodes after the food source was changed at 3 days of age (Fig. 4A); however, the brood size was the same as that of the control nematodes (Fig. 4B). The longevity effects due to FC were unlikely to be attributed to the so-called dietary restriction, because the EPSes appeared to have longevity effects on the nematodes fed OP (Fig. 1G).

**Fig. 4A.**
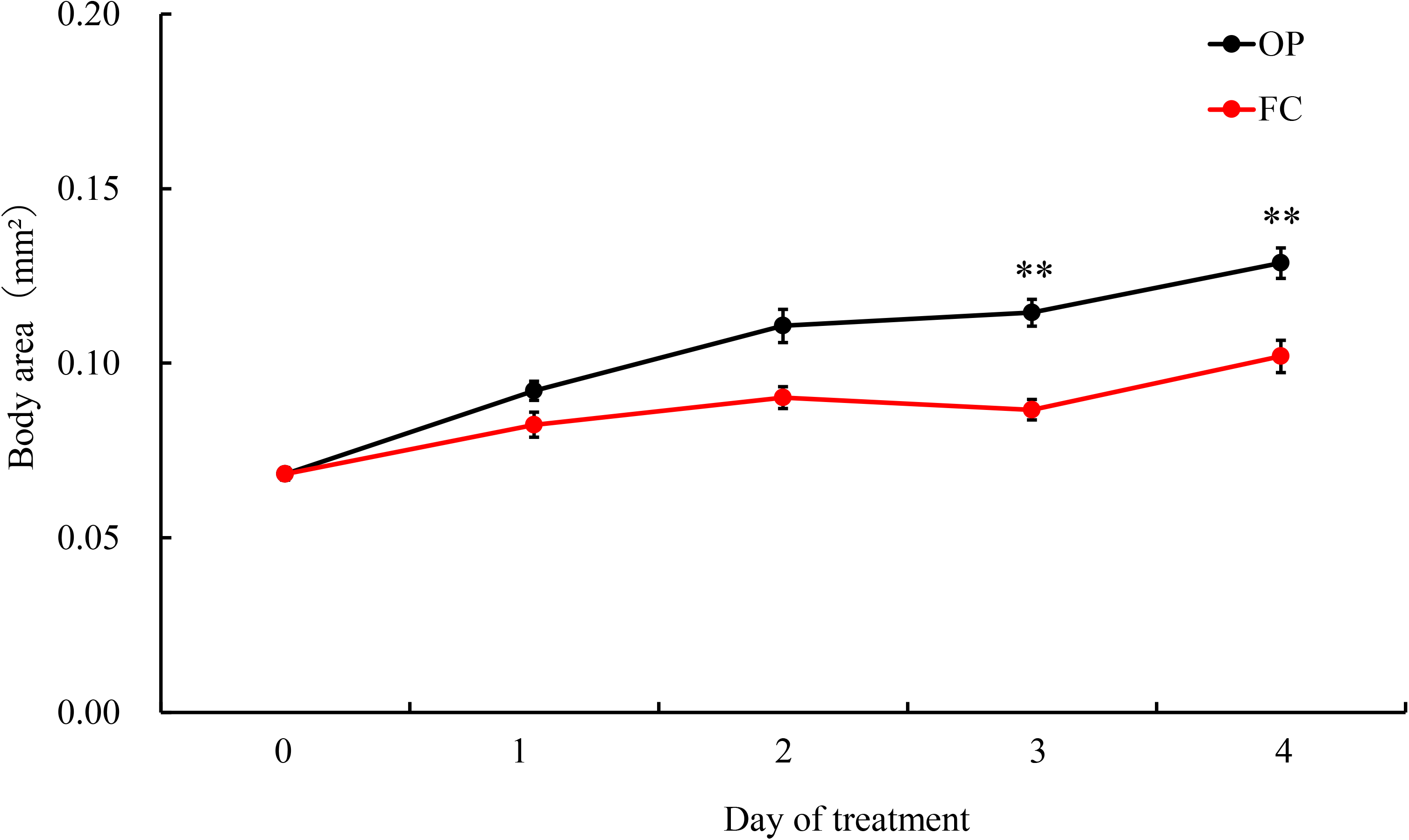
Nutritional value of FC for worms. Growth curve of worms fed OP or aerobically grown FC. Images of adult nematodes were captured using a digital microscope, and the area of the worm’s projection was measured and used as an index of the body size. The body sizes of the worms fed aerobically grown FC was less than that of the control worms fed only OP. All results are shown as means ± standard error. Asterisks indicate significant differences (** *p* < 0.01) from control worms fed OP. The assay was performed with 10 worms and repeated twice.

**Fig. 4B.**
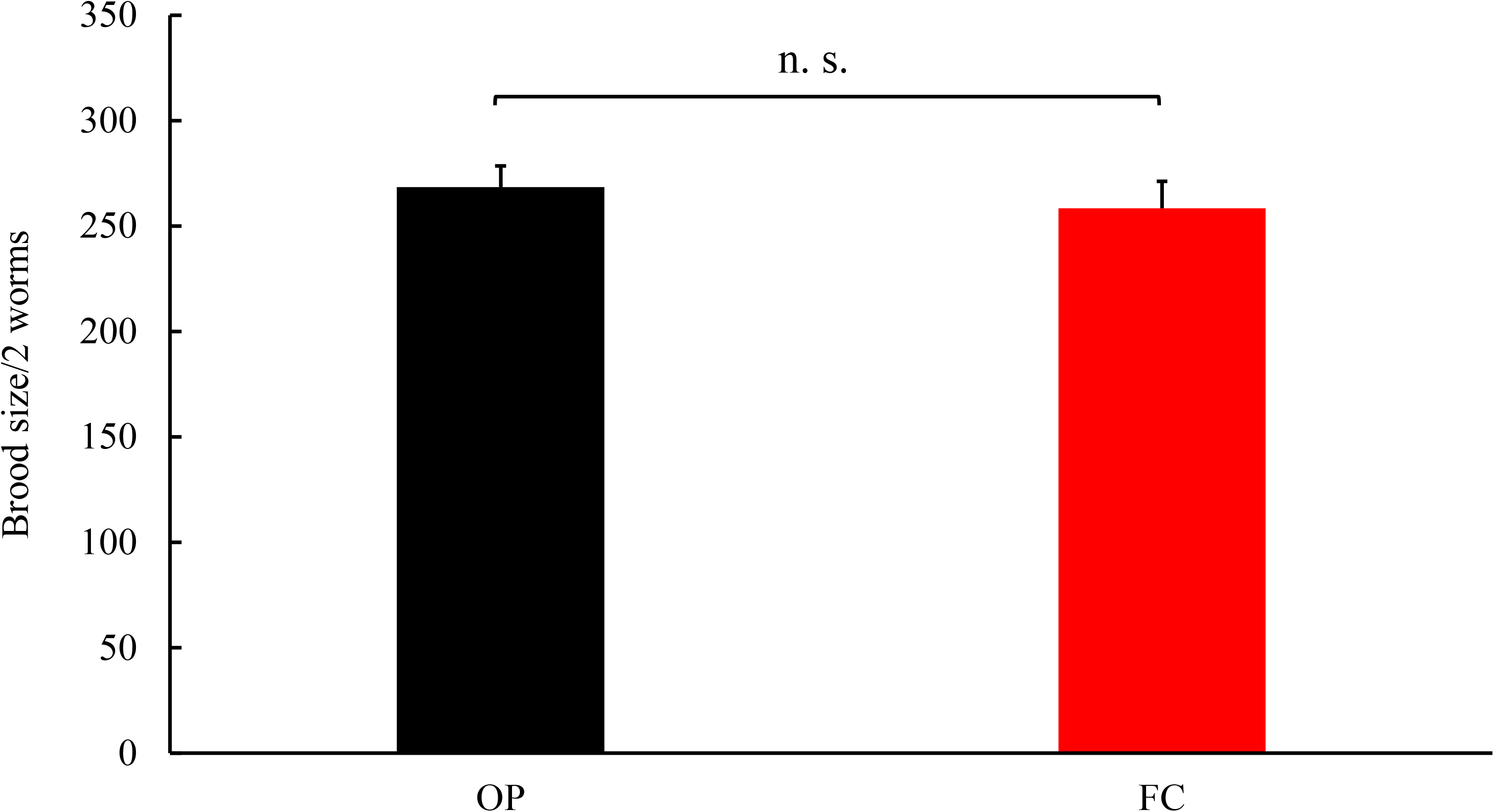
Nutritional value of FC for worms. Brood size of worms fed aerobically grown FC. Brood size of FC-fed worms did not vary from that of control worms fed OP. The total brood size was determined for more than 12 worms. Error bars represent the standard error. Each assay was repeated thrice.

The mechanism of the anti-senescence effects of FC which prolonged the lifespan and enhanced the host defense of nematodes was investigated by examining mutants lacking defense-related signaling pathways. DAF-16, the mammalian homolog of the FOXO transcription factor, is located at the end of the insulin/insulin-like growth factor signaling pathway (IGF-1) and is activated to enhance host resistance when upstream IGF-1 signals decrease due to conditions such as dietary restriction. Another transcription factor SKN-1 at the end of the p38 MAPK (PMK-1) pathway regulates the expression of xenobiotic detoxification genes (26). When mutant nematodes with defects in host defense-associated genes (*daf-16* and *skn-1*) were fed FC, the bacteria extended the lifespan of *daf-16* and *pmk-1* mutants, but not that of *skn-1* mutants (Fig. 5A). Although the cognitive ability of *daf-16* mutants (Fig. 5B) was lower than that of wild-type N2 (Fig. 2B), FC-fed mutants showed higher chemotaxis index than the control fed OP at the age of 7 days (Fig. 5B). However, FC failed to improve the cognitive ability of *skn-1* mutants (Fig. 5C). Furthermore, the survival time of *skn-1* mutants after infection with *S.* Enteritidis or *S*. *aureus* was similar between OP and FC feeding (Fig 5D): the *skn-1* mutants lost tolerance to both the pathogens.

**Fig. 5A.**
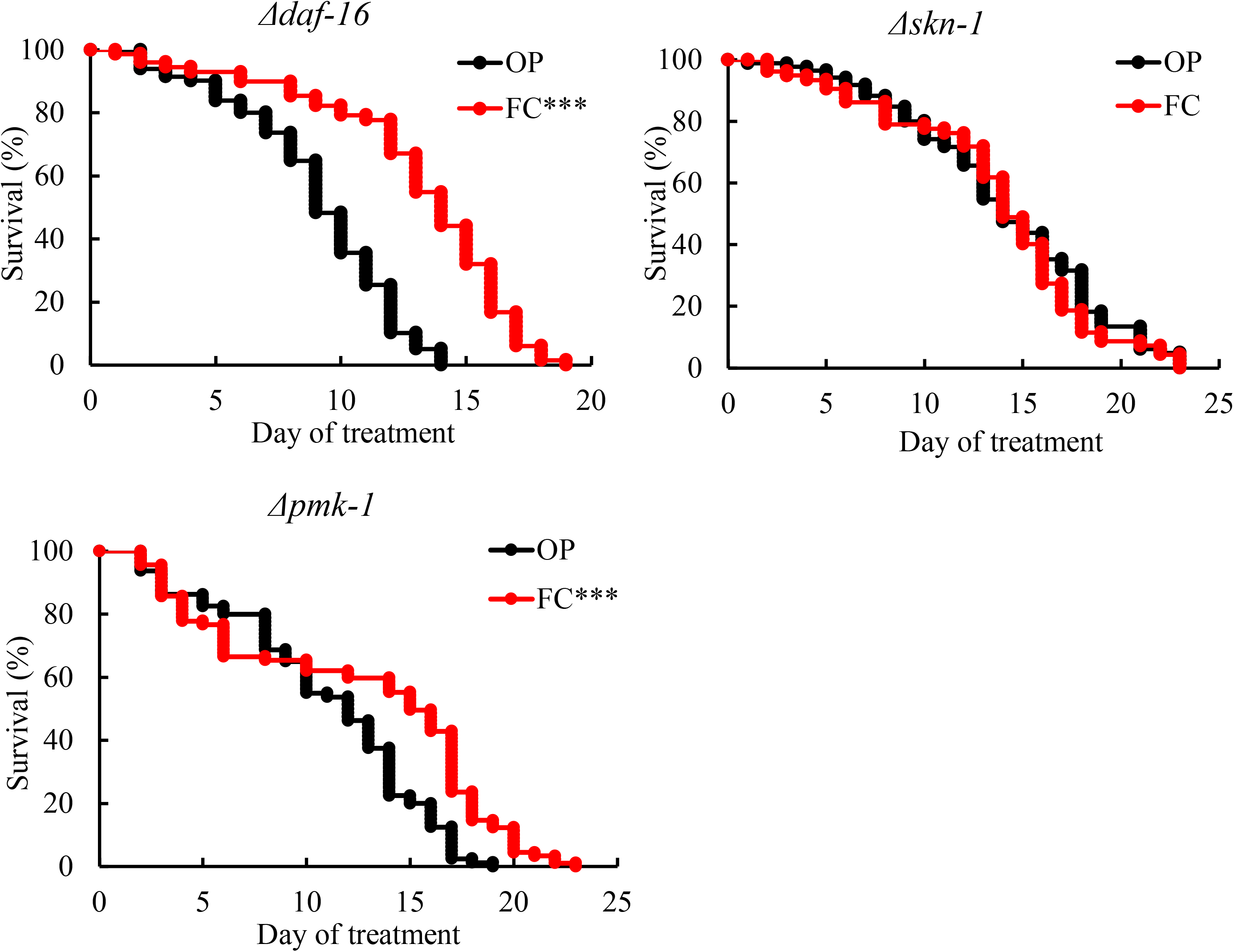
Effects of FC on *C. elegans* mutants. Survival curves of *daf-16 (mu86)* and *skn-1 (zu67)* mutant hermaphrodites fed with or without FC from 3 days of age. The worms were 3 days of age at day 0. FC extended the lifespan of the *daf-16* and *pmk-1* mutants, but not of *skn-1* mutant. Each assay was repeated twice by using aerobically cultured organisms and once with anaerobically cultured bacteria with *daf-16*; thrice by using aerobically cultured organisms and twice with anaerobically cultured cells with *skn-1*; and four times by using aerobically cultured organisms with *pmk-1* mutants. Table 2 summarizes the data obtained from all experiments, and the curves were drawn based on representative experiments. Asterisks indicate statistically significant differences (*** *p* < 0.001) compared to control worms fed OP.

**Fig. 5B.**
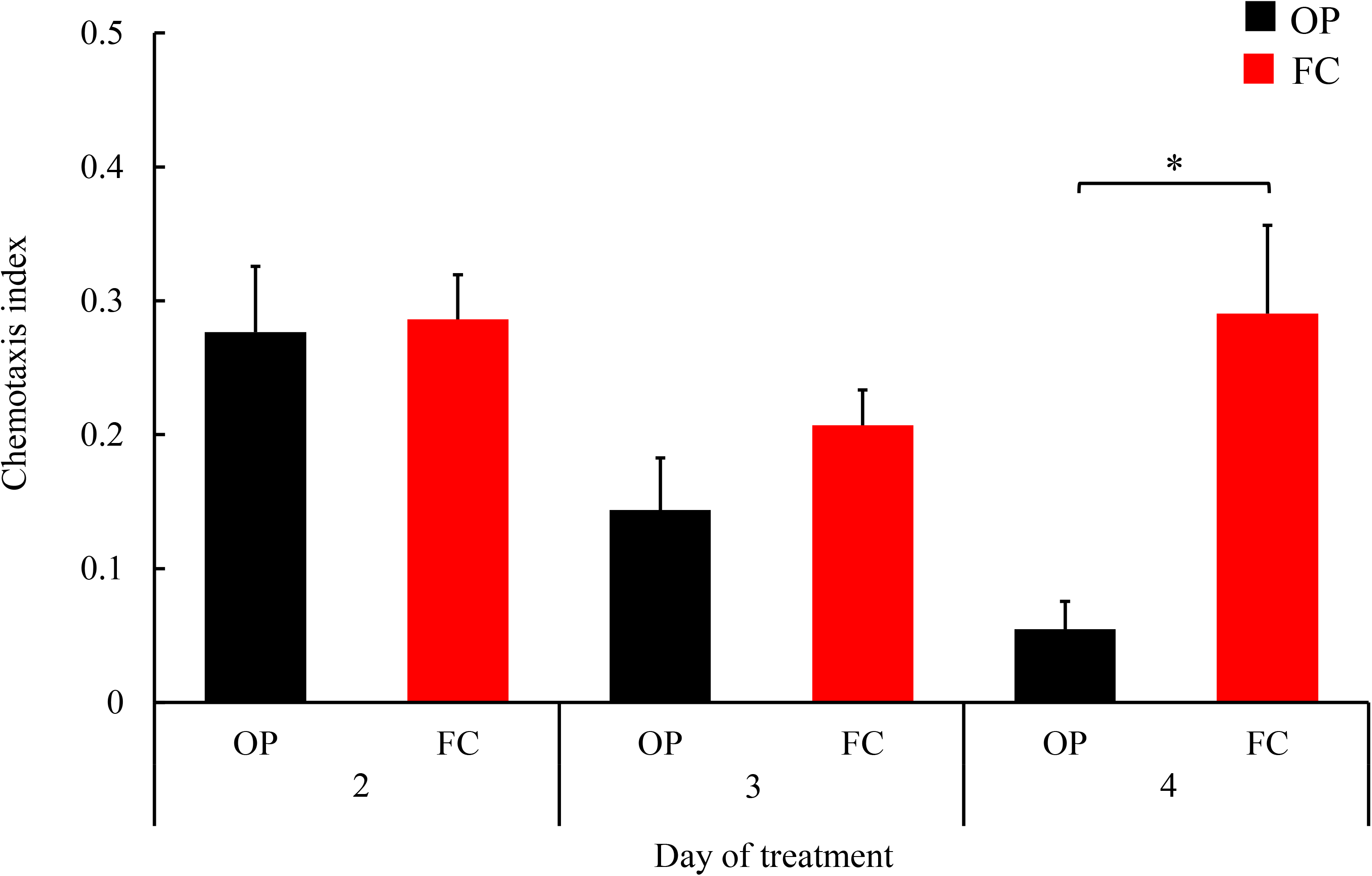
Effects of FC on *C. elegans* mutants. The chemotaxis index (CI) of *Δdaf-16* fed OP or aerobically grown FC from 3 days of age. FC-fed mutants showed higher CI than the control fed OP at 7 days of age. Each bar represents the average values of a minimum of 200 worms. Error bars represent the standard error. The assay was performed in triplicate and repeated three times. Asterisks indicate a statistically significant difference (* *p* < 0.05) compared to control worms fed OP.

**Fig. 5C.**
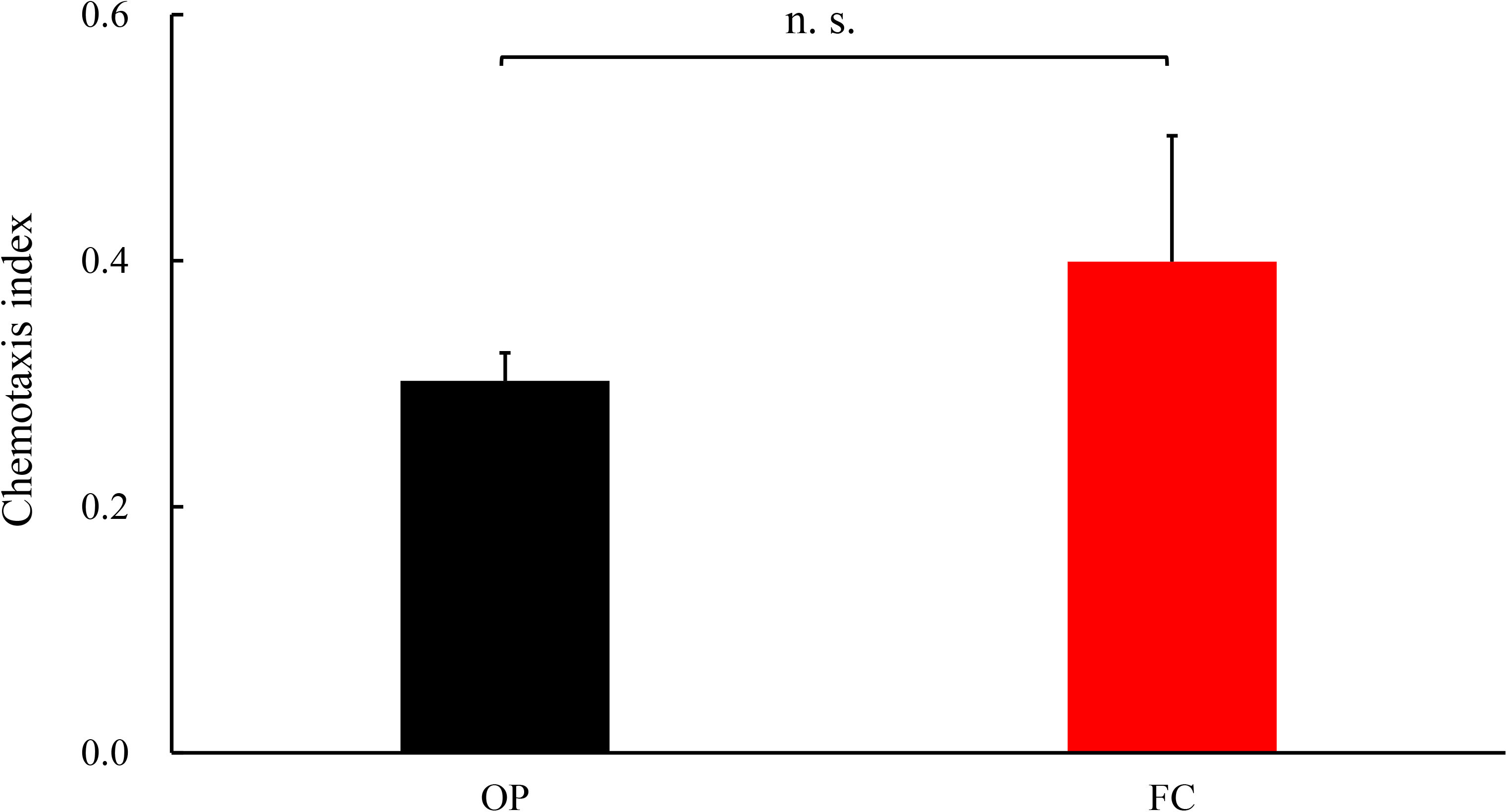
Effects of FC on *C. elegans* mutants. The CI of the 7-day-old *Δskn-1* fed OP or aerobically grown FC from 3 days of age. The *skn-1* mutants fed FC failed to show improved cognition. The assay was repeated three times. Each bar represents the average values of a minimum of 200 worms.

**Fig. 5D.**
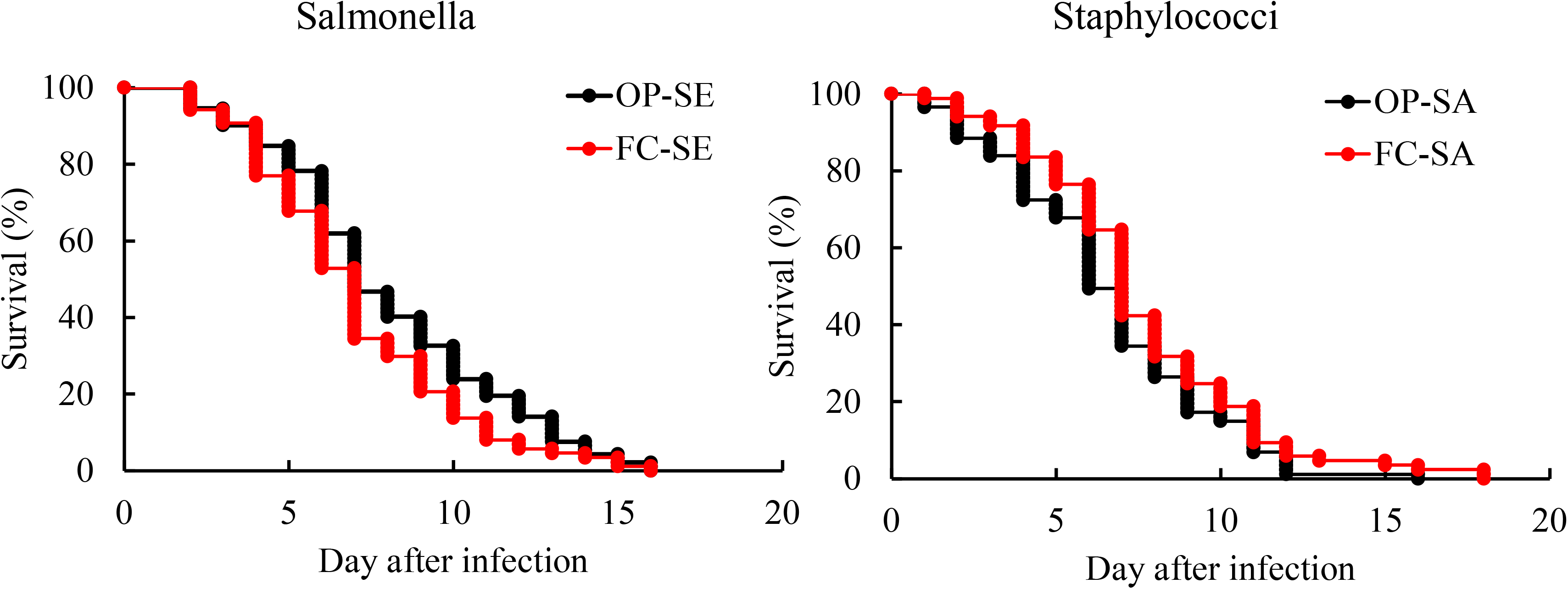
Effects of FC on *C. elegans* mutants. Effects of FC on the resistance of *C. elegans* mutant to *Salmonella enterica* serovar Enteritidis (SE) or *Staphylococcus aureus* (SA). The *skn-1* mutant fed OP for 3 days after hatching were transferred to the diet of aerobically grown FC or OP as a control. After 4 days, the nematodes were transferred to SE or SA plates, and survival curves were drawn. Survival times after infections with SE or SA were similar between OP- and FC-fed *skn-1* mutants. Each assay was performed once using SE or SA. Table 3 summarizes the data obtained from all experiments, and the curves were drawn on the basis of representative experiments.

As SKN-1 is similar to nuclear erythroid 2-related factor 2 (Nrf2) in mammals, we investigated whether FC could enhance the Nrf2 antioxidant response pathway in mammalian cells. We measured the transcription of the HO-1 gene, which is regulated by Nrf2. The mRNA level of HO-1 increased by more than 14 times after inoculation with FC cells, compared to that in non-inoculated macrophages (Fig. 6). However, heat-killed bacteria could not produce this effect. The EPS could induce the transcription of HO-1 in a dose-dependent manner. Although the effects of EPS were not statistically significant, it worked synergistically with heat-killed FC cells. Thus, EPS can also be implicated as an effector that causes beneficial effects *in vivo*.

**Fig. 6.**
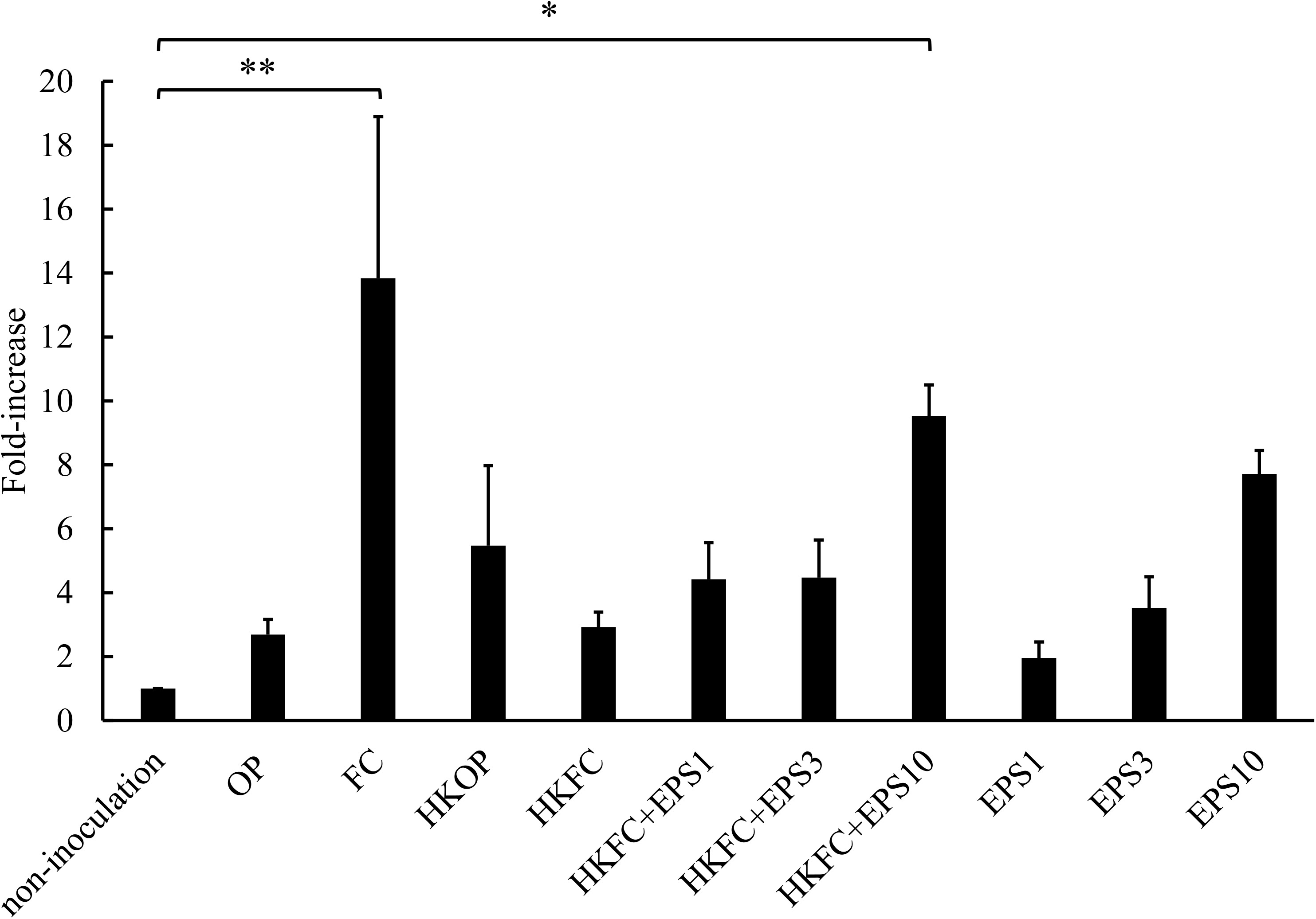
The HO-1 mRNA expression in J774.1 cells treated with FC. Cells were pretreated with viable FC or OP, heat-killed FC (HKFC) or heat-killed OP (HKOP), and with/without exopolysaccharide (EPS) (1, 3, or 10 mg/mL) for 24 h. The mRNA of HO-1 increased after inoculation with FC or HKFC + EPS10 (EPS concentration is 10 mg/mL) cells, compared to non-inoculated cells. Data represent the mean ± standard error of three independent experiments. Asterisks indicate significant differences (* *p* < 0.05, ** *p* < 0.01) from non-inoculated cells. The assay was repeated three times.

## DISCUSSION

In this study, we showed that *L. lactis* subsp. *cremoris* strain FC extended the lifespan and health span of nematodes without affecting the brood size. Interestingly, FC made the nematodes resilient to infections and stressors, decreased AGEs levels, delayed senescence and improved cognitive ability. The increased autofluorescence from AGEs and deterioration of cognitive ability were alleviated in either 9- or 7-day-old nematodes, respectively; unlike the conventional lifespan assay, these examinations rapidly determine whether probiotic bacteria and any chemicals exert longevity effects in *C. elegans*.

A previous study suggested that the decline in chemotaxis index could be attributed to sarcopenia, but not impaired cognition (27). However, in this study, comparatively young adult worms (3- to 7-day-old) were used, with reportedly well-maintained integrity of their nervous system and muscles at this age (28). Indeed, in the chemotaxis assay, the number of worms moving out of the center from where they were settled first was similar between 5- and 7-day-old worms; thus, locomotive function was not significantly impaired in 7-day-old worms. Furthermore, the response to odorants is low in worms aged more than 9 or 11 days due to functional deterioration of neuromuscular signaling (28). Thus, the decrease in the perception of the attractant might be responsible for the low chemotaxis index in older worms. Even in mice, audibility, another sensory neuron function, is lost earlier than the other symptoms of senescence (21).

Noteworthy, even the international food standard OP can become pathogenic in old nematodes. Worms fed heat-killed OP lived longer than those fed live OP; the longevity afforded by FC was possibly attributed to their harmless nature. However, the nematodes fed FC lived longer than those fed heat-killed OP. Thus, the longevity by FC might not be simply attributed to its non-pathogenic property.

The EPS of the FC also extended the lifespan of *C. elegans*. Donato et al. recently reported that extracellular polymetric substances produced *in vivo* by *Bacillus subtilis* were implicated in the prolonged longevity of nematodes (29). In this study, FC was recovered alive from the nematodes (Suppl. Fig. 1), and the nematodes fed living FC had better survival than those fed heat-killed FC. Thus, we assumed that the EPS synthesized *in vivo* might contribute to the longevity effect of living FC. This assumption for FC differs from what we observed previously in *Bifidobacterium infantis*, whereby both the cell wall or protoplast alone could also extend the lifespan of worms in the same manner as the living bifidobacteria (30). This implies that the active constituents and mechanism of action by each probiotic bacterium differ between species.

Although the brood size of nematodes fed FC was normal, their body size and lipid amount were less than those of nematodes fed OP. FC, being a low-calorie food, is assumed to prolong the lifespan through the so-called caloric restriction. However, even the administration of EPS could extend the lifespan of worms fed OP. Hence, dietary (calorie) restriction alone does not explain the longevity effect afforded by FC. Furthermore, caloric restriction reportedly extends the lifespan of worms mainly via DAF-16(31, 32), and Huang et al. showed that the longevity afforded by aspirin resulted in decreased fat storage via DAF-16 (33). In contrast, Xu et al. indicated that *C. elegans* mutants of microtubule regulators which are involved in both neuronal function and neuronal structure maintenance, could achieve longevity via increased fat storage in a DAF-16-dependent manner (34). After all, mechanistic alterations in both lipid production and break down may be more critical effectors than the resultant fat storage or reduction in aging. In this study, FC-fed worms showed reduction in fat storage, but the effects were independent of DAF-16. Because of these controversial findings, whether decreased fat content is associated with the longevity of FC-fed worms is not yet clear.

Even under stresses such as exposure to heat or juglone and infections with *S.* Enteritidis or *S*. *aureus*, nematodes fed FC had better survival than those fed OP. This resilience seems to be attributed to tolerance rather than resistance, because the number of pathogens recovered was similar between infected nematodes fed FC and those fed OP. This is likely associated with the better epithelial barrier function in the worms fed FC, particularly *via* zonula occludens protein-1, which was indicated by less leakage in the intestinal barrier function assay (Smurf assay) (35).

The helix-loop-helix protein 30 (HLH-30) transcription factor (an ortholog of TFEB) is reportedly translocated to the nucleus during infection with staphylococci and regulates the transcriptional immune response (36) by affecting the expression of genes involved in well-established pathways for host defense, such as PMK-1, antimicrobial genes (lysozymes and C-type lectins), and autophagy genes. Of note, transcription factor SKN-1 at the end of the PMK-1 pathway might also be activated *via* bacterial signals, lipids, XBP-1, and PMK-1, and can be suppressed by ASI neurons, TOR signaling, DAF-2, GSK-3, WDR-23, and some miRNAs (37–41). Nevertheless, HLH-30 likely plays an important role in the immune response of *C. elegans* to *S. aureus via* the upregulation of autophagy genes. However, *C. elegans* has not yet been reported as a general model of intracellular infection (42). Since autophagy is expected to be a defense system for intracellular pathogens, the contribution of autophagy to the tolerance induced by FC was not expected. However, Zhao et al. suggested that bacteria can enter the pharyngeal tissue, particularly, the pharyngeal muscle near the grinder, through the damaged pharyngeal cuticle in 4-day-old *C. elegans* and remain latent from early-to mid-adulthood. The infection then recrudesces in later life when the worm’s intrinsic defense deteriorates with senescence (25). Autophagy seems to play an important role in the production of intracellular bacteria (43), and worms that fail to control bacteria die with a median age of 12 days owing to pharyngeal swelling (25). In contrast, worms that survive until mid-adulthood seem to live longer and die due to marked atrophy of the pharyngeal posterior bulb.

Hence, the interventions for longevity of *C. elegans* need to target at the early death due to a swollen pharynx or later death due to an atrophied pharynx. FC enhanced host defense against infection, heat, and juglone, while prolonging the mean lifespan. The prolonged longevity effects of FC could be attributed to the alleviation of pharyngeal swelling and atrophy by host defense *via* SKN-1. Zhao et al. reported that a 42-fold greater number of bacteria was recovered from swollen pharynxes than from normal ones. However, in this study, no difference was noted between the number of bacteria recovered from the whole bodies of FC-fed worms and the control, because the bacterial population in pharynxes seems to account only for a minor percentage of the total bacteria present in the whole body.

For the longevity due to FC, SKN-1 was pivotal, whereas DAF-16, a well-known transcription factor that regulates genes promoting stress resistance and longevity (44), was not. SKN-1 belongs to the cap-n-collar family (CNC) of basic leucine zipper transcription factors and regulates the expression of xenobiotic detoxification genes (26). Reduction of the blue autofluorescence, an index for AGEs (22), in the worms fed FC might also be dependent on SKN-1 (45–47). The mammalian transcription factor Nrf2 of the CNC family is homologous to SKN-1. In this study, FC and the EPS caused higher HO-1 transcription in mouse macrophages than in non-treated cells. Thus, FC may modulate immunological responses and ameliorate senescence in both nematodes and mammal cells.

In conclusion, by using *C. elegans*, established as an economic model host to select probiotic candidates in our previous studies, we showed that *L. lactis* subsp. *cremoris* and the EPS could provide anti-senescence effects by activating host defense *via* the transcription factor SKN-1 in nematodes and by activation of Nrf2 in murine macrophages.

## MATERIALS AND METHODS

### Nematode

*C. elegans* Bristol strain N2 and its derivative mutant strains were kindly provided by the Caenorhabditis Genetics Center, University of Minnesota. The following mutants were used in this study: KU25 *pmk-1 (km25)*, EU1 *skn-1 (zu67)*, and CF1038 *daf-16 (mu86)*. Nematodes were maintained and propagated on nematode growth medium (NGM), according to standard techniques (48). OP was used as the standard feed for nematode cultivation and was grown on tryptone soy agar (Nissui Pharmaceutical, Tokyo, Japan). Cultured bacteria (100 mg wet weight) were suspended in 0.5 mL M9 buffer, and 50 μL of the resulting bacterial suspension was then spread on peptone-free modified NGM (mNGM) in 5.0-cm-diameter plates to feed the worms.

### Bacterial strains

FC (a stock strain of Fujicco Co., Kobe, Japan) was used as a test food source for *C. elegans* and was anaerobically cultured in M17 broth (Becton Dickinson, Franklin Lakes, NJ, USA) with 0.5 % lactose and M17 agar (Becton Dickinson) with 0.5 % lactose at 25°C for 48 h. *Salmonella enterica* serovar Enteritidis strain SE1, originally isolated from a diarrheal specimen, was used as a gram-negative pathogen. *Staphylococcus aureus* strain 96-55-17A was used as a gram-positive pathogen. These pathogenic bacteria were reported as infection models by using *C. elegans* (49) and were grown on tryptone soya agar at 37 °C for 24 h.

### Determination of *C. elegans* life span

Eggs were recovered from adult *C. elegans* worms after exposure to sodium hypochlorite/sodium hydroxide solution, as previously described (30). The egg suspension was incubated overnight at 25°C to allow hatching, and the resulting suspension of L1 stage worms was centrifuged at 156 × *g* for 1 min. After the supernatant was removed by aspiration, the remaining larvae were transferred onto fresh mNGM plates covered with OP and incubated at 25°C. Pubescence was synchronized by allowing the worms to feed on OP for 2 days until maturation, as the reproductive system is known to regulate aging in *C. elegans* (50). Nematocidal assays were performed by adding 3-day-old adult worms to each mNGM plate covered with lawns of FC. The plates were incubated at 25°C, and the numbers of live and dead worms were scored every 24 h. At 25°C, the worms produce progeny that develop into adults within 3 days; therefore, identifying the original worms is difficult. Overestimation of the number of living worms was avoided by transferring the original worms daily to fresh mNGM plates for 4 days until they completed their egg-laying phase at 7 days of age. The worms were then transferred to fresh mNGM plates every alternate day. A worm was considered dead when it failed to respond to a gentle touch with a worm picker. Worms that died resulting from adhering to the wall of the plate were not included in the analysis. Nematocidal assays are generally performed using NGM agar plates containing peptone, which facilitates the proliferation of the overlaid bacteria. However, it is reported that the composition of NGM influences the virulence of bacteria, and that the *in situ* production of metabolites by bacteria growing on the medium could also be fatal to nematodes (51). Thus, the possibility of bacterial-induced nematocidal effects from nutrients in the medium was avoided by performing the nematocidal assays on mNGM plates lacking peptone. Heat-killed bacteria were prepared at 100°C for 10 min. Each assay was performed in duplicate and repeated more than twice, unless otherwise stated.

The mean lifespan (MLS) was estimated using the following formula (52):

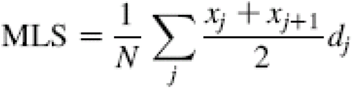

where *d_j_* is the number of worms that died in the age interval (x_j_, x_j_ + 1), and *N* is the total number of worms. The standard error (SE) of the estimated MLS was calculated using the following equation:

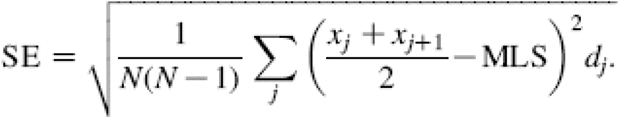

The maximum lifespan was calculated as the MLS of the top 15% longest-living worms in each group.

### Locomotory scoring of ageing nematodes

The motility of worms at different ages was examined using a scoring method described previously (30, 53). In brief, worms were classified as class “A” when they showed spontaneous movement or vigorous locomotion in response to prodding; class ‘‘B’’ when they did not move unless prodded or appeared to have uncoordinated movement; and class ‘‘C’’ when they moved only their head and/or tail in response to prodding. Dead worms were classified as class “D.” A minimum of 60 worms fed OP or FC were scored.

### Measurement of autofluorescence as an aging marker in *C. elegans*

We recently developed a method to measure autofluorescence as an aging marker for individual worms (22). In brief, 9-day-old worms fed OP or FC from the age of 3 days were washed with M9 buffer and placed into 1.0 µL of M9 buffer on a 384-well black plate (Stem, Tokyo, Japan) covered with Saran Wrap (Asahi Kasei, Tokyo, Japan). The blank (M9 buffer only) data were checked three times for each well, considering the fluctuation across wells. Minimal detection limits and quantifiable limits were determined on the basis of blank data on each day as μ (mean of the blank) + 3.29σ (standard deviation) and μ + √2 × 10σ, respectively. The autofluorescence in the worm body was captured using a multimode grating microplate reader model SH-9000Lab (excitation, 340 nm; emission, 430 nm; Corona Electric, Ibaraki, Japan). After measurement, each worm was individually maintained on a 4-cm-diameter plate covered with OP or FC (2 mg/10 μL M9 buffer) at 25°C. The assay was performed with more than 20 worms and was repeated three times.

### Measurement of body size

Adult worm size was determined as established previously (5). Three-day-old adult worms were placed on mNGM plates covered with lawns of FC. The plates were incubated at 25°C, and the body size of live worms was measured every 24 h until they reached 7 days of age. Images of adult nematodes were captured using an STZ-161 microscope (Shimadzu, Kyoto, Japan) and a USB camera L-835 (Hozan, Osaka, Japan). Body size was analyzed using Adobe Photoshop elements and ImageJ version 1.52a. software developed by the National Institutes of Health. In this system, the area of a worm’s projection was estimated automatically and used as the index of body size.

### Lipid staining in *C. elegans*

Staining was performed as previously described with minor modifications (54). In brief, 7-day-old worms fed OP or FC for 4 days were washed three times with M9 buffer. The samples were fixed in 50% isopropanol in PBS for 15 min on ice. Oil red O stock solution (0.5 g/100 mL in isopropanol; Sigma-Aldrich, St. Louis, MO, USA) was diluted in distilled water (dH_2_O) to 60% working solution and filtered using 0.2 μm membrane. Fixed worms were incubated in working solution at 25°C for 20 min. Stained worms were washed with M9 buffer containing 0.5% Triton X-100 and mounted onto glass slides by using M9 buffer + 0.5% Triton X-100 for imaging by using an Olympus BX53 equipped with a color camera DP74 (Olympus, Tokyo, Japan). The measured dye values were normalized to density values determined for the body size, as described above. Next, the dye values were measured using ImageQuant TL version 8 (GE Healthcare, Chicago, IL, USA) and Adobe Photoshop Elements were normalized to density values per mm^2^ of a worm’s projection area. The assay was performed twice.

### Preparation of exopolysaccharides from FC

The EPSes were isolated from fermented milk incubated at 26°C for 8 h with FC, as described previously (12). Trichloroacetic acid (TCA) was added at 4% w/v to fermented milk to remove the precipitate and bacterial cells, and the resulting suspension was centrifuged at 18,890 × *g* for 20 min at 4°C. EPS was precipitated from the supernatant with 1.5 mL of chilled ethanol, collected using a spatula, dissolved in dH_2_O water, and precipitated at 18,890 × *g* for 20 min at 4°C with 4% TCA. The EPS was obtained by dialysis of the supernatant against dH_2_O water for 3 days and subsequent lyophilization.

EPS was administered in the form of a liposome so that *C. elegans* could ingest hydrophilic substances (24). The liposome was prepared and administered as previously described, with minor modifications (24). In brief, the EPS was dissolved in dH_2_O water (15 mg/mL). Next, L-α-phosphatidylcholine was added to the solution at 40 mg/mL, and liposomes were produced by mixing the solution at 65°C and passing through a Nucleopore track-etched membrane (pore size, 5.0 µm; Whatman, Newton, MA, USA) by using a Mini-Extruder (Avanti Polar Lipids, Alabaster, AL, USA). For comparison, liposomes containing dH_2_O were used as a control. A nematode was fed 25 μL OP (400 mg/mL) and 25 μL liposomes containing EPS on mNGM from 3 days of age.

### Chemotaxis assays

This assay was performed as described previously (55), with some modification. Synchronized wild-type worms, *skn-1*, and *daf-16* mutants were fed OP or FC from 3 days of age. The 7-day-old worms were collected and assayed on 90 mm mNGM plates. Next, 1 µL 1.0 M sodium azide along with 1 µL odorant 0.1% benzaldehyde in 100% ethanol was dropped onto the center of attractant regions A and D (2 cm in diameter) on the plates. At the center of regions B and C located at the opposite end of the attractant regions, 1 µL 100% ethanol as control and 1 µL sodium azide were dropped. Next, about 50–70 worms washed with M9 buffer were immediately transferred to the center of the plate; only worms that moved spontaneously and showed vigorous locomotion in response to prodding were used. The assay plates were incubated at 25°C for 2 h, and then the number of worms in each quadrant was scored. The chemotaxis index (CI) was calculated using the formula: CI = (number of worms in both attractant quadrants – number of worms in both control quadrants)/total number of worms on the assay plate. Worms that adhered to the plate wall were not included in the analysis. Each assay was performed in triplicate and repeated three times.

### Influence of feed on resistance against pathogenic bacterial infection

Resistance against pathogenic bacterial infections was determined as previously described (5). After hatching, the nematodes were fed OP for 3 days, and then the worms were assigned to either a control group that was continued to be fed OP or to a group that was fed FC for 4 days. The 7-day-old worms were then transferred onto *S.* Enteritidis or *S. aureus* lawns. Each group was incubated at 25°C. The numbers of live and dead worms were scored every 24 h. Survival rates were compared between the groups of worms grown on different feeds before the *S.* Enteritidis or *S. aureus* infection. Each assay was performed in duplicate and repeated three times.

### Measurement of the number of bacteria in nematodes

The number of bacterial cells in the nematodes was determined according to a previous method (56) with some modifications. The surface bacteria were killed by immersing more than five worms in 500 μL gentamycin solution (1 mg/mL) for 30 min in a 0.5 mL microtube. After the worms were washed five times in M9 buffer, each nematode was placed in a microtube containing 50 µL M9 buffer and mechanically disrupted using a Mini Cordless Grinder (Funakoshi, Tokyo, Japan). The volume was adjusted to 1 mL by using M9 buffer. Before the culture on agar, the homogenate solution was diluted in M9 buffer, which was plated on tryptone soya agar for OP, *S.* Enteritidis, and *S. aureus* at 37°C, or on M17 with lactose agar for FC at 25°C. All plates were incubated for 48 h.

### Smurf assay for intestinal barrier function

Worms were infected following the infection method described above. Infected or non-infected worms (12 day old) on mNGM agar were transferred to S-basal medium containing heat-killed OP mixed with acid blue 9 as a blue food dye (5% w/v in S-basal solution with cholesterol; Tokyo Chemical Industry Co., Tokyo, Japan) at 25°C for 3 h. Worms were washed in M9 buffer for five times before they were anesthetized in M9 buffer containing 500 mM sodium azide. Subsequently, the worms were observed on the agar pad for the presence of blue dye in the body cavity by using an Axiovert 135M microscope (Carl Zeiss, Jena, Germany) and USB camera L-835 (Hozan).

### Stress resistance assays

In this method, 7-day-old worms fed OP or FC from 3 days of age were exposed to heat and various oxidative stresses. Thermal tolerance was assessed by incubating worms at 35°C for 7 h, and the survival rate was calculated after the worms were recovered on mNGM seeded with OP or FC at 25°C for 24 h. An oxidative stress assay was conducted by transferring 7-day-old worms onto mNGM containing 250 µM juglone (a reactive oxygen species generator) for 2h, and viability was scored after a 15 h recovery period on normal mNGM seeded with OP or FC. Juglone was dissolved in 100% ethanol, and the assays were performed according to the method described previously (57, 58) with some modification. For other oxidative stress assays, worms were placed onto mNGM containing 1.0 mM paraquat or in 24-well tissue culture plates containing 7.0 or 9.0 mM cupric chloride added to K-medium (53 mM NaCl, 32 mM KCl) (59), and the numbers of live and dead worms were either scored every hour or daily, respectively. The survival of worms was determined using touch-provoked movements. Worms were scored as dead when the animals failed to respond to mechanical stimuli with a worm picker. The assay was performed on 100 worms and was repeated three times.

### Brood size

Brood size was determined as previously established (30). Eggs isolated with a sodium hypochlorite or sodium hydroxide solution were allowed to develop up to 3 days of age on mNGM plates coated with OP at 25°C. Two hermaphrodites were selected and transferred to an mNGM plate covered with a lawn of OP or FC. The parental animals were transferred every 24 h to fresh mNGM plates until the end of the reproductive period. The resulting progeny were left to develop for 3 days, and the progeny number was then determined. Each assay was performed with five plates and was repeated three times.

### Cell culture and mRNA expression

Initially, the Hep G2 cell line (BPS Bioscience, San Diego, CA, USA) containing a luciferase gene under the control of an antioxidant response element was used. Although FC was assumed to activate the antioxidant response element of this reporter cell line, it failed unexpectedly, potentially due to poor bacterial ingestion by hepatic cells. Thus, the murine macrophage cell line J774.1 (JCRB0018; JCRB Cell Bank, Osaka, Japan) was used instead. These cells were grown in RPMI-1640 (Wako, Osaka, Japan), supplemented with 0.5 % v/v StemSure 10 mmol/l 2-mercaptoethanol solution (Wako) and 10% v/v fetal bovine serum (#F7524; Sigma-Aldrich), as the cell medium. Cells were grown to approximately 80% confluence in 25 cm^2^ polystyrene tissue culture flasks at 37°C in a 5% CO_2_ incubator. Before bacterial treatment, the cells were transferred to a 24-well plate (5 × 10^5^ cells/500 µL cell medium/well; Thermo Fisher Scientific, Waltham, MA, USA) overnight. The bacteria were collected in tubes and washed with PBS. For heat-killed bacteria, bacteria were boiled at 100°C for 10 min and then adjusted to 100 mg/mL with cell medium. The J774.1 cells were cultured with living or heat-killed bacteria of either OP or FC (0.5 mg/500 μL cell medium) for 24 h. For EPS, cells were treated alone (EPS: 1, 3, or 10 mg/mL) or mixed with heat-killed FC (0.5 mg/500 μL cell medium). Total mRNA was extracted from either the bacteria and/or EPS-treated cells, or untreated cells by using the NucleoSpin RNA kit (Macherey-Nagel, Duren, Germany), according to manufacturer’s instructions. cDNA was synthesized using ReverTra Ace qPCR RT Master Mix with gDNA remover (TOYOBO, Osaka, Japan). The transcription levels of the HO-1 gene were detected using the Luna Universal qPCR Master Mix (New England Biolabs Japan, Tokyo, Japan). Real-time PCR was performed using Mastercycler ep realplex^2^ (Eppendorf, Hamburg, Germany). Information on the primers used is shown in Table 1. Target mRNA expression was normalized to the expression of a reference mRNA (actb, encoding the housekeeping protein β-actin), and the fold change was calculated on the basis of the threshold cycle (ΔΔCT) method (60).

### Statistical analysis

Nematode survival was calculated using the Kaplan–Meier method, and survival differences were tested for significance by using the log-rank test. Autofluorescence values were analyzed using the Mann–Whitney *U* test. Oil red O strain values, brood size, and chemotaxis assays for the *skn-1* mutant were compared using Student’s *t*-test. Body size and chemotaxis assays for wild-type N2 and *daf-16* mutants and the number of bacterial cells in *C. elegans* were analyzed using a two-factor factorial ANOVA and Tukey–Kramer test or Scheffe’s *F* test for multiple comparisons. The chi-squared test was used to assess the heat stress and oxidative stress induced by juglone. The mRNA expression of the cells was analyzed using single-factor ANOVA and Dunnett test. Where significance was observed, data were classified as * *p* < 0.05, ** *p* < 0.01, and *** *p* < 0.001. All statistical analyses were performed using Microsoft Excel supplemented with the add-in software + Statcel 3 and 4 (OMS, Tokyo, Japan).

## ACKNOWLEDGMENTS

The nematodes used in this study were kindly provided by the Caenorhabditis Genetics Center, which is funded by the NIH Office of Research Infrastructure Programs (P40 OD010440). This study was supported in part by the Institute for Fermentation (No. G-2020-3-057; Osaka, Japan) to T. K. We would like to thank Editage (www.editage.com) for English language editing.

---

Supplementary Fig. 1. Longitudinal changes in the numbers of FC and OP cells recovered from nematodes. FC was recovered alive from nematodes. The result is presented as means ± standard errors of the means.

