## Supplemental Figure 1 for "Prolonged Lifespan, Ameliorated Cognition, and Improved Host Defense of *Caenorhabditis elegans* by *Lactococcus lactis* subsp. *cremoris*"

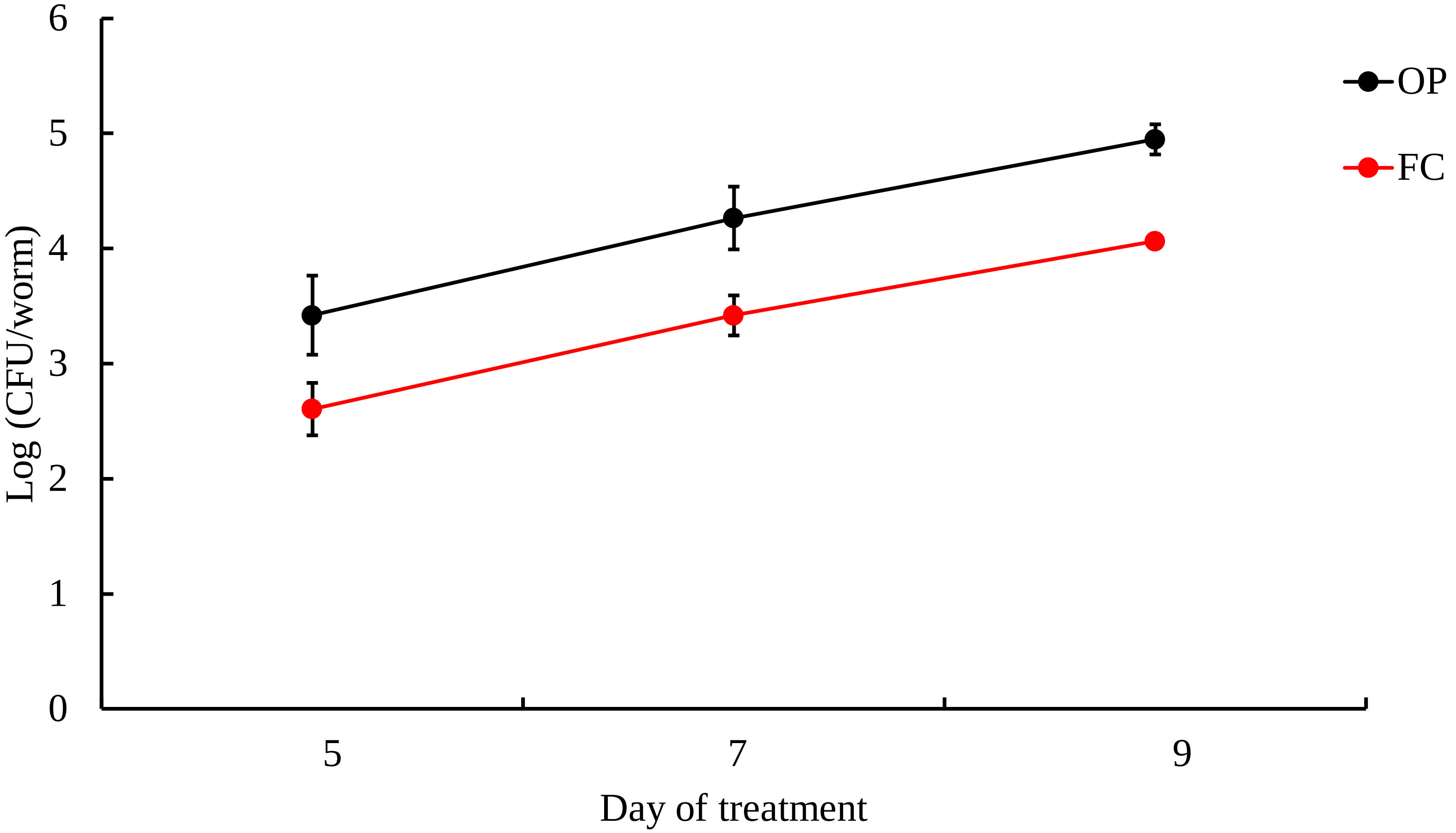


Supplementary Fig. 1. Longitudinal changes in the numbers of FC and OP cells recovered from nematodes. FC was recovered alive from nematodes. The result is presented as means ± standard errors of the means.
